# The cell cycle controls centrosome growth by regulating the dynamics of two distinct scaffolds

**DOI:** 10.64898/2026.11.20.768102

**Authors:** Siu-Shing Wong, Zachary M. Wilmott, Emma L. Roberts, Kwai-Yin Chau, Thomas L. Steinacker, Jordan W. Raff

**Affiliations:** The Sir William Dunn School of Pathology, University of Oxford; Oxford, OX1 3RE, UK; Department of Biochemistry, University of Oxford; Oxford, OX1 3QU, UK; Spark Data Systems Ltd, Hotwells, Bristol; BS8 4UQ, UK

## Abstract

Centrosomes comprise a pair of centrioles that organise a surrounding pericentriolar material (PCM) that usually grows as cells prepare to enter mitosis, stops growing during mitosis, and then shrinks as cells exit mitosis. In most models of centrosome growth, the PCM stops growing because cells run out of a limiting component. Using experiments and mathematical modelling in the early *Drosophila* embryo, we show that the embryonic cell cycle oscillator (CCO) can generate these centrosome growth kinetics without relying on the depletion of a limiting component. At the start of each cycle centrioles assemble two dynamic and interconnected PCM scaffolds comprising Spd-2/CEP192 and Cnn/CDK5RAP2. As embryos approach mitosis, rising CCO activity destabilises the Spd-2 scaffold, slowing centrosome growth, but stabilises the Cnn scaffold, “locking” it into a low-dynamic state. At the end of mitosis CCO activity drops, triggering rapid Cnn scaffold disassembly. Thus, the CCO drives centrosome growth dynamics by regulating the stability of two interconnected PCM scaffolds.

## Introduction

Cells contain many different organelles that perform a myriad of functions. A newborn proliferating cell will have to double the amount of each organelle prior to the next round of cell division, but how growth is regulated to ensure that cells make the right amount of each organelle at the right time and place is poorly understood (1–3).

Centrosomes are membraneless organelles that function as major microtubule-organising centres in many animal cells (4–7). They are an interesting model for studying organelle growth as most cells are born with just a single centrosome that duplicates precisely once every cell cycle. Centrosomes comprise a central pair of mother/daughter centrioles that recruit a matrix of pericentriolar material (PCM) around themselves (Fig. 1A). In interphase, centrioles usually organise a small amount of PCM that is tightly associated with the mother centrioles (8–11). The centrosomes duplicate in S-phase and, as cells prepare to enter mitosis, the PCM starts to grow around the mother centrioles in a process termed centrosome maturation (12). This allows the two mitotic centrosomes to organise more microtubules (MTs) to form the poles of the bipolar mitotic spindle, upon which the duplicated chromosomes will segregate. The mitotic PCM stops growing during mitosis (13, 14) and is then rapidly disassembled as cells exit mitosis (15–17).

**Fig. 1.**
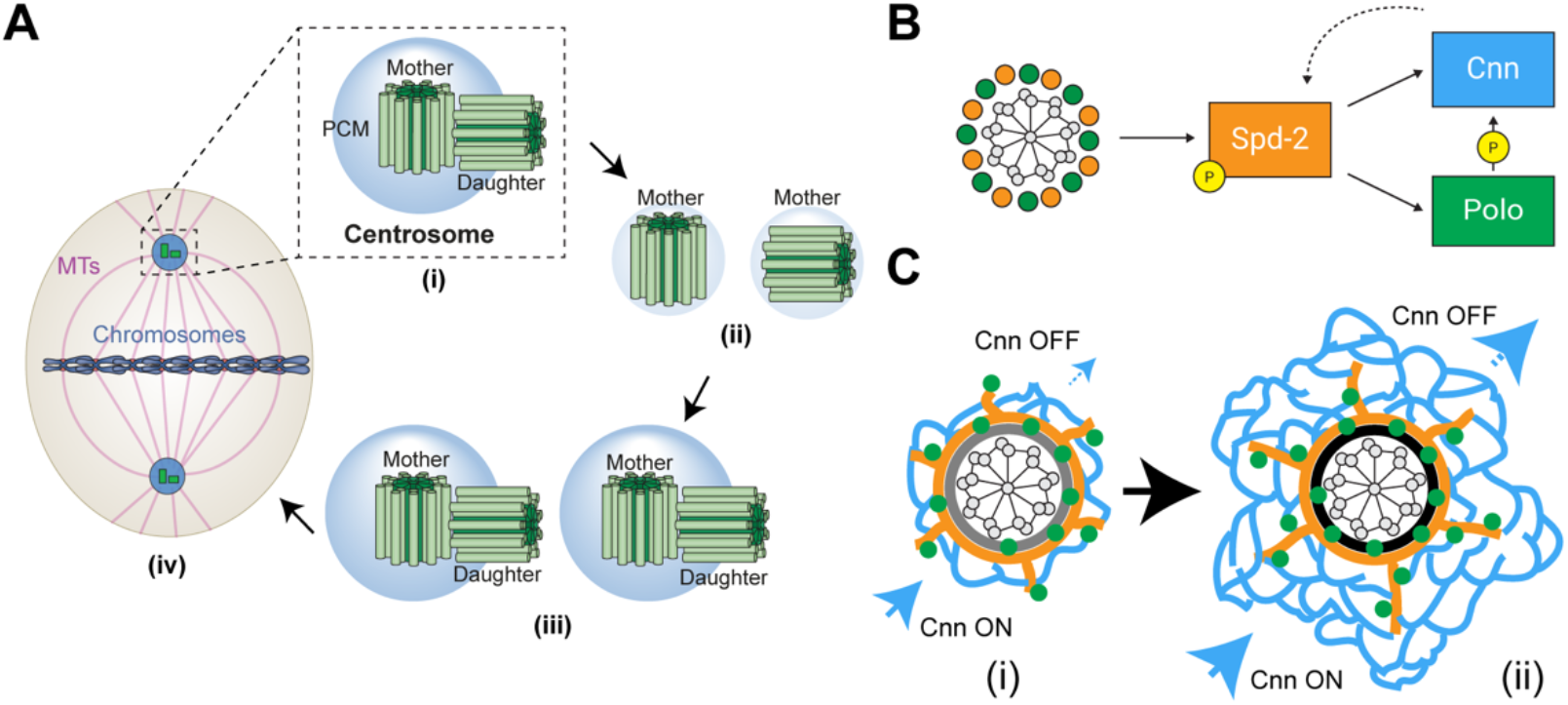
The centrosome cycle and mitotic PCM growth. **(A)** The centrosome cycle. (i) A mitotic centrosome comprises a mother and a daughter centriole surrounded by pericentriolar material (PCM). (ii) As cells exit mitosis, most PCM is disassembled and the two centrioles separate, so each now acts as a mother. (iii) In S-phase, each mother centriole builds a new daughter. As cells get ready to enter mitosis, the PCM grows around the mother centrioles. (iv) In mitosis, the two centrosomes organise microtubules (MTs) to form the poles of the bipolar spindle. **(B)** Schematic summary of mitotic PCM scaffold assembly in *Drosophila* embryos. Polo (*green*) and Spd-2 (*orange*) are concentrated at centrioles throughout the cell cycle. As embryos prepare to enter mitosis (which in the rapidly dividing early *Drosophila* embryo, which has no G1 or G2 phases, occurs as soon as the embryos exit the previous mitosis), centrioles generate a pulse of Polo activity that phosphorylates Spd-2. This allows Spd-2 to form a weak scaffold that fluxes away from the centrioles recruiting more Polo and also Cnn (*blue*). The Polo bound to the Spd-2 scaffold phosphorylates Cnn, driving Cnn scaffold assembly. The Cnn scaffold cannot recruit Spd-2 (so generation of the Spd-2 scaffold remains dependent on the centrioles), but it helps stabilise the Spd-2 scaffold (indicated by the dashed arrow). **(C)** Schematic illustrates how the Cnn scaffold initially grows (i) because its ON rate is larger than its OFF rate. As the scaffold enlarges, its OFF rate increases, so it can grow until it reaches a steady state (ii), where its ON rate is balanced by its OFF rate.

Polo/PLK1 kinase is the major driver of mitotic centrosome growth in many systems (18, 19). In *Drosophila* embryos, Polo (Plk1/PLK-1 in humans/worms) phosphorylates Spd-2 (CEP192/SPD-2 in humans/worms) at centrioles as embryos prepare to enter mitosis (Fig. 1B) (20, 21). This allows Spd-2 to form a weak scaffold that can flux away from the centriole, recruiting an additional pool of Polo that can phosphorylate the large coiled-coil protein Cnn (CDK5RAP2/SPD-5 in humans/worms) to promote the assembly of a large scaffolding structure around the mother centriole (22–26) (Fig. 1C). This scaffold provides mechanical strength and helps to recruit other proteins to the expanding mitotic PCM. There is strong evidence that the worm homologues of these proteins function in a similar manner (27–34), and this also appears to be the case for the human homologues (35–39), although additional proteins such as Pericentrin may also have an important role (40).

Recent studies in fly embryos have shown that the dynamics of mitotic centrosome assembly are surprisingly complicated (21, 41). Centrosomal levels of Polo and Spd-2 rise and fall during the assembly process, but Cnn levels continue to rise and then plateau. A mathematical model in which a pulse of Polo activity at the centriole initiates the Polo/Spd-2/Cnn interactions described above was proposed to explain these dynamics (21). In this model, centrosomes stop growing during mitosis because the Cnn scaffold reaches a steady-state size—where Cnn incorporation (Cnn ON) driven by the Spd-2 scaffold is balanced by Cnn loss at the periphery (Cnn OFF) (Fig. 1C).

Two other mathematical models have been proposed to explain the kinetics of centrosome growth in early worm and/or fly embryos: a model based on the assumption that centrosomes are autocatalytic droplets formed by liquid-liquid phase separation (LLPS) (42) and a model based on the assumption that catalytic growth of the centrosome is promoted by a shared enzyme pool (43). In both models, centrosomes stop growing during mitosis because a limiting component becomes depleted from the cytoplasm. This limiting component hypothesis is based on the observation that SPD-2/CEP192 behaves as such component in the early worm embryo (29). This hypothesis has been influential in the centrosome and organelle size control fields (2, 44–47).

Interestingly, recent data indicated that the cell cycle oscillator (CCO) has a role in terminating centrosome growth in early *Drosophila* embryos (41), a possibility not considered in any of the previous models. Here we examine how the CCO influences the kinetics of mitotic centrosome scaffold assembly in this system. Using FRAP, fluorescence correlation spectroscopy (FCS) and mathematical modelling, we provide evidence that Spd-2 and Cnn are not significantly depleted from the cytoplasm during centrosome assembly in the *Drosophila* embryo. Instead, we find that the CCO dynamically regulates the rate at which Spd-2/CEP192 and Cnn/CDK5RAP2 are recruited to, and disassemble from, the mitotic PCM during each nuclear cycle. At the start of each cycle, the centrioles start to grow by assembling dynamic Spd-2 and Cnn scaffolds. As the embryo prepares to enter mitosis, however, increasing CCO activity destabilises the Spd-2 scaffold—so the growth of the Cnn scaffold slows—but stabilises the existing Cnn scaffold. Thus, *Drosophila* embryos enter mitosis with a shrinking Spd-2 scaffold and a static Cnn scaffold. At the end of mitosis, the rapid decrease in CCO activity stimulates the rapid disassembly of the Cnn scaffold.

We conclude that the cell cycle regulated assembly of a dynamic Spd-2/CEP192 scaffold and a more stable Cnn/CDK5RAP2 scaffold can explain the kinetics of centrosome growth in the early fly embryo without a requirement to invoke a limiting component. This model may provide a paradigm for understanding mitotic centrosome growth more generally.

## Results

### Measuring the recruitment dynamics of Spd-2 and Cnn through the cell cycle

We wanted to measure how Spd-2 and Cnn dynamics changed during the cell cycle in syncytial *Drosophila* embryos, where we can observe multiple centrosomes progressing through multiple **nuclear cycles (NCs)** in near-synchrony. To do this, we used fluorescence recovery after photobleaching (FRAP) in embryos expressing CRISPR knock-in NeonGreen (NG) fusions of either Spd-2 (Spd-2-NG) or Cnn (NG-Cnn), photobleaching a small sample of centrosomes at 2-minute intervals as embryos progressed through NC12 (Fig. 2A, Video S1, 2). Because NC length varies slightly between embryos, we rescaled time in each embryo between centrosome separation, which marks the approximate start of S-phase (t=0.0), and nuclear envelope breakdown (NEB), which marks the approximate start of mitosis (t=1.0). This allowed FRAP measurements from centrosomes in different embryos to be compared at approximately equivalent cell-cycle stages (Fig. 2B, S1A).

**Fig. 2.**
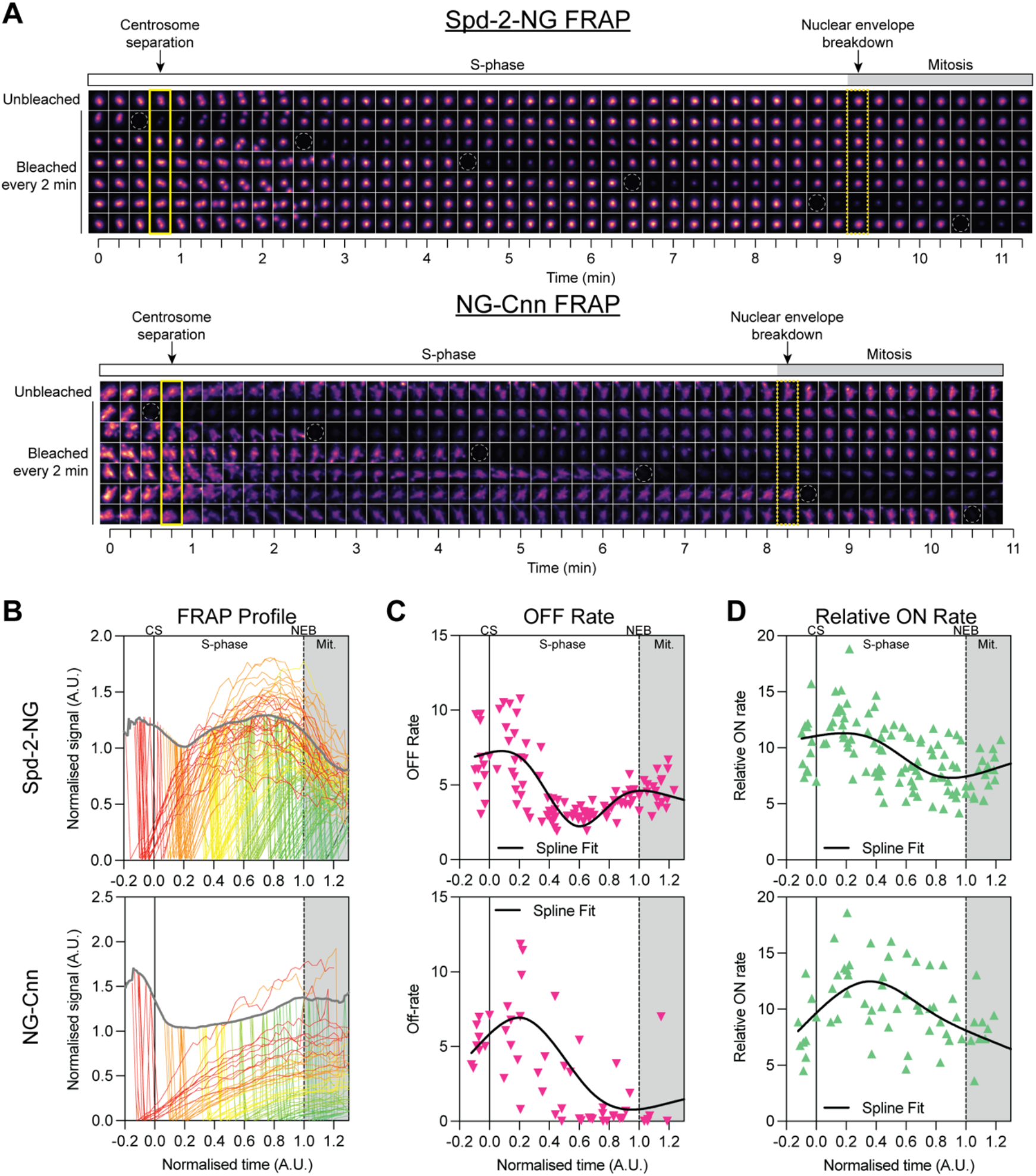
Spd-2 and Cnn scaffold dynamics diverge as embryos approach mitosis. **(A)** Stills from time-lapse videos of NC12 embryos expressing Spd-2-NG (top) or NG-Cnn (bottom). Each row initially shows a separating centrosome pair (note that the centrosomes can be observed to separate at slightly different times). The centrosomes shown were then photobleached at staggered 2-min intervals as the embryo proceeded through NC12. White dotted circles mark the bleaching timepoints. Solid and dashed yellow boxes mark the approximate time of centrosome separation (CS) and nuclear envelope breakdown (NEB), respectively. **(B)** FRAP recovery curves for Spd-2-NG (top) and NG-Cnn (bottom), plotted against normalised time (0 = CS, 1 = NEB; grey shading, mitosis). Each coloured line shows the average fluorescence recovery of 3–4 centrosomes that were bleached at the same timepoint; lines are colour-coded by the bleaching timepoint (e.g., centrosomes bleached at t=0.0 are shown in *red*; note that because time is normalised, the centrosomes bleached at each timepoint are spread out slightly over normalised time). The thick *grey* line shows the mean signal of the unbleached centrosomes. **(C, D)** Graphs plot how the OFF (C) and ON (D) rates of Spd-2 (top) and Cnn (bottom) vary over NC12. Each point is calculated from one FRAP curve shown in (B). Black lines are spline fits showing the trend across the cycle. CS, centrosome separation; NEB, nuclear envelope breakdown; Mit., mitosis. N = 12–20 embryos; n > 500 centrosomes.

Interpreting these FRAP experiments is complicated by the fact that total centrosomal Spd-2 and Cnn levels are not constant during the recovery period—they increase or decrease as the centrosomes progress through the cycle. This means the raw FRAP recovery curves reflect the exchange of bleached for unbleached protein, as well as the underlying rise or fall in the total amount of protein at the centrosome. To isolate the exchange component, we subtracted the FRAP curve, *y*_*F*_, from its corresponding unbleached control growth curve, *y*, and normalised the result. This difference curve was well described by an exponential decay to a non-zero plateau, *y* − *y*_*F*_ = *j* + (1 − *j*)*e*^−*k*_off_(*t*−*t*_0_)^ (Fig. S1B) (see *Appendix I* for mathematical derivation). The plateau, *j*, represents the fraction of protein that doesn’t exchange on the timescale of observation. The decay constant, *k*_off_, is the probability that a bound molecule dissociates per unit time (Fig. 2C).

We also calculated a *relative* ON rate (Fig. 2D). At the photobleaching timepoint, *t*_*F*_, the rate at which new, fluorescent material is incorporated is given by 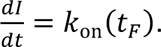 Because *I*(*t*) is measured in arbitrary units, however, this yields an ON rate in arbitrary units. Converting this to an absolute incorporation rate would require knowledge of absolute molecule numbers, which we could not determine. Hence, while the OFF rate we measure is an absolute rate constant, the ON rate is only known on a relative scale.

This analysis revealed that the Spd-2 and Cnn OFF rates were relatively high and rising as centrosomes started to assemble at the start of S-phase of NC12 but soon started to decline as S-phase progressed (Fig. 2C). The Spd-2 OFF rate then started to rise, whereas the Cnn OFF rate continued downwards and was very low by the time the embryo entered mitosis. The relative Spd-2 and Cnn ON rates broadly exhibited a similar pattern to their OFF rates, although, unlike the Cnn OFF rate, the Cnn ON rate never approached zero (Fig. 2D). We conclude that the Spd-2 and Cnn scaffolds are both relatively dynamic as centrosomes start to assemble in early S-phase (exhibiting relatively high ON and OFF rates), become less dynamic as the cycle progresses, but then diverge in their behaviour as the embryos prepare to enter mitosis: the Spd-2 scaffold becomes more dynamic, whereas the Cnn scaffold enters a low dynamic state—exhibiting low ON and very low OFF rates.

### The embryo cytoplasm does not appear to be depleted of Spd-2 or Cnn during centrosome assembly

As discussed above, it has long been postulated that centrosomes stop growing in mitosis because a limiting component (SPD-2 in worm embryos) becomes depleted from the cytoplasm during the assembly process (29, 42–44). To test if this might contribute to the centrosome growth dynamics we observe in fly embryos, we used FCS to monitor how the concentration of Spd-2-NG and NG-Cnn (expressed as CRISPR knock-ins from their endogenous loci) changed during NC12 (Fig. 3). The cytoplasmic concentration of Spd-2 remained relatively constant throughout S-phase as centrosomes assembled, while for Cnn it fell slightly (*yellow boxes*, Fig. 3). The cytoplasmic concentration of both proteins then rose slightly as embryos entered mitosis. We conclude that these scaffold building blocks are not (Spd-2), or are only modestly (Cnn), depleted from the cytoplasm during the assembly process. Thus, the decline of the Spd-2 scaffold and the plateauing of the Cnn scaffold as the embryos enter mitosis is not due to the cytoplasmic depletion of either protein.

**Fig. 3.**
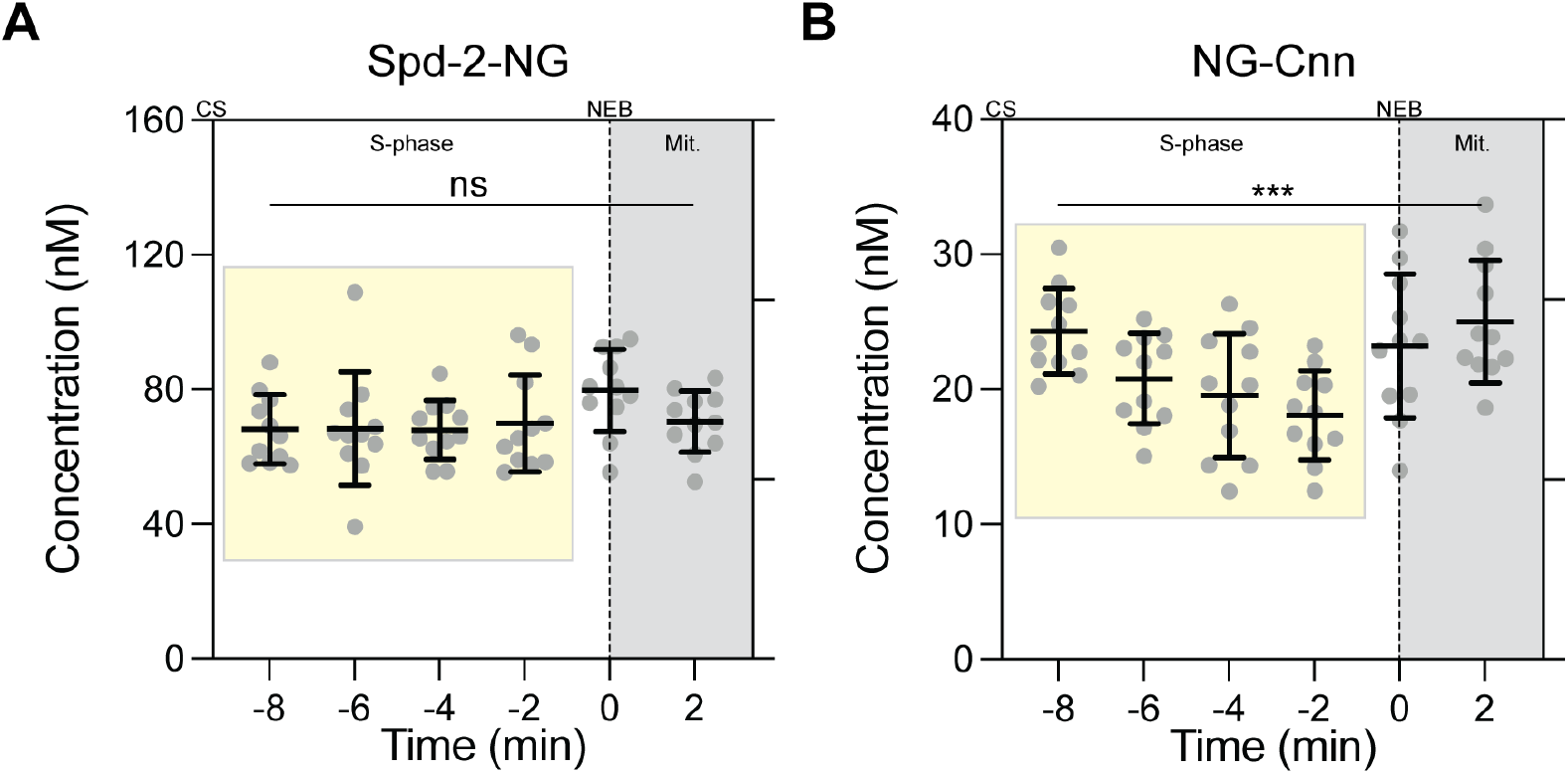
Centrosome growth does not substantially deplete cytoplasmic Spd-2 or Cnn. **(A, B)** Cytoplasmic concentrations of Spd-2-NG (A) and NG-Cnn (B), measured by FCS at 2min intervals during NC12. Time is relative to nuclear envelope breakdown (NEB; t = 0, dashed line). *Yellow* boxes mark S-phase and *grey* shading marks mitosis. Each point is one embryo (the mean of 4–6 × 10s FCS recordings). Bars show mean ± SD. N = 12-15 embryos per timepoint. Statistical significance was tested by a nonparametric Friedman test (***P < 0.001; ns, not significant).

### A model of mitotic centrosome growth that incorporates regulation by the CCO

Two important assumptions of our previous model of mitotic PCM assembly (21)—that the OFF rate of the Spd-2 scaffold remains constant during assembly and that the OFF rate of the Cnn scaffold increases as its size increases (Fig. 1B,C)—are not captured by the turnover dynamics we measure here. The other mathematical models of mitotic centrosome assembly also do not explicitly incorporate the strong cell-cycle dependence of scaffold turnover revealed here (42, 43). We therefore developed a model of mitotic PCM scaffold assembly that incorporates how Spd-2 and Cnn dynamics change during the cell cycle.

This new model remains based on the reciprocal interactions that are known to be important for Polo/PLK1-stimulated Spd-2/CEP192 and Cnn/CDK5RAP2 scaffold assembly in fly embryos (Fig. 4A). Centrosome growth is initiated at the start of each cycle by a pulse of centriolar Polo activity, which is likely triggered by activating the centriole protein Ana1/CEP295 to recruit Polo (21). This centriolar pulse of Polo promotes the recruitment of cytoplasmic Spd-2 into a centrosomal scaffold that can flux outwards from the centriole, recruiting more Polo (20, 23). The pool of Polo bound to the Spd-2 scaffold in turn promotes the recruitment of cytoplasmic Cnn into a second centrosomal scaffold. For simplicity, we reduce the recruitment of Cnn by Spd-2 via Polo into a single step in which the initial Polo-associated Spd-2 scaffold recruits Cnn. The Cnn scaffold cannot recruit new Spd-2 molecules, but it can stabilise the Spd-2 scaffold (23). In this way, the two scaffold layers are reciprocally coupled: the Spd-2 scaffold directly promotes Cnn scaffold assembly, and the Cnn scaffold stabilises the Spd-2 scaffold, thereby indirectly promoting further Spd-2 and Cnn scaffold growth (*red circle*, Fig. 4A).

**Fig. 4.**
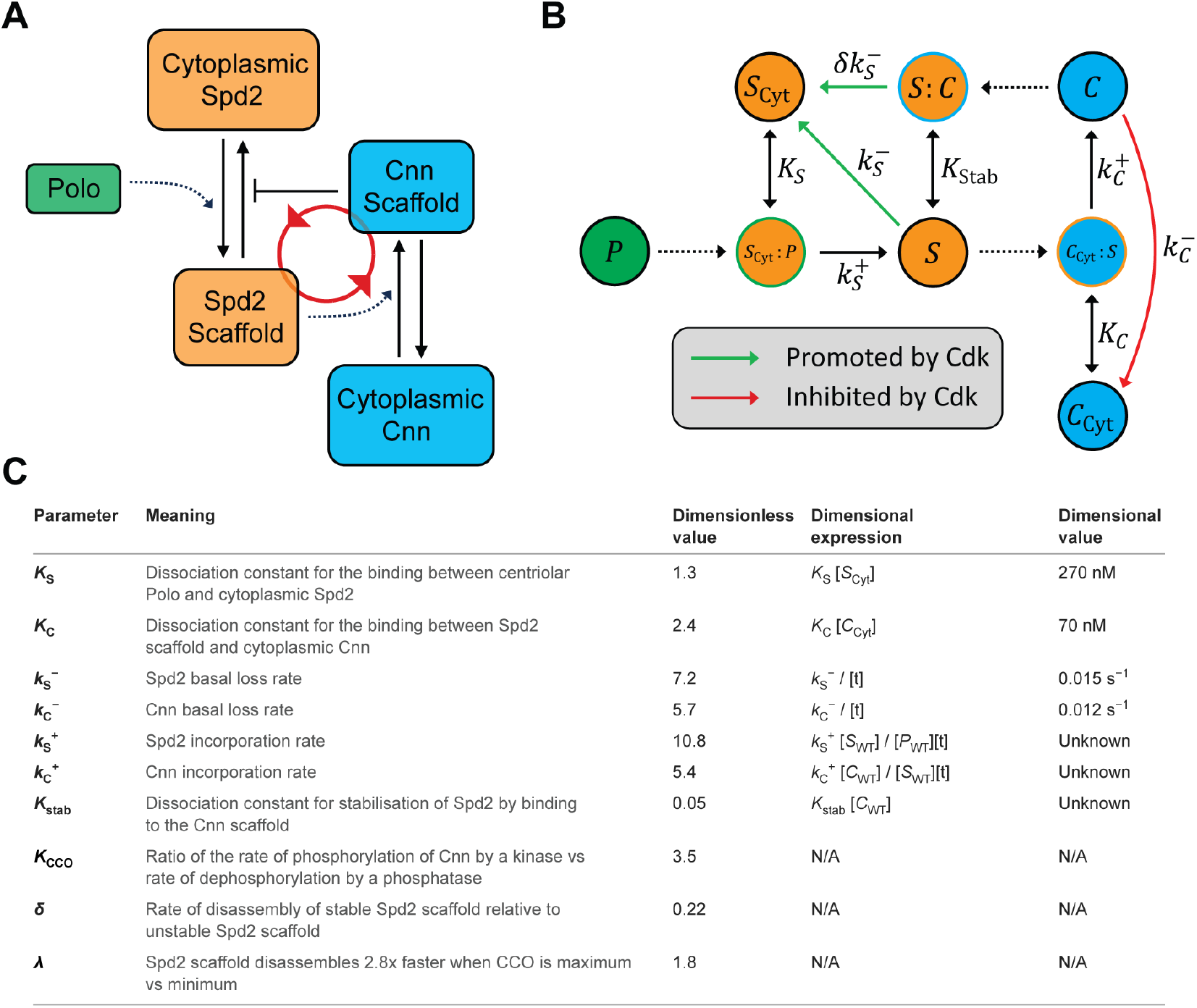
A model in which the cell cycle oscillator regulates the assembly of coupled Spd-2 and Cnn scaffolds. **(A)** Schematic overview of the model. A pulse of Polo activity at the centriole drives recruitment of cytoplasmic Spd-2 into a scaffold (dotted arrow). The Spd-2 scaffold then drives recruitment of cytoplasmic Cnn into a second scaffold (dotted arrow). For simplicity, the Polo that Spd-2 recruits is not modelled as a separate step. The Cnn scaffold stabilises the Spd-2 scaffold by slowing its disassembly (blunt arrow). This two-way coupling forms a positive feedback loop (*red* arrows). **(B)** Reaction scheme of the model (based on Equations 1 and 2). **(C)** Table summarises the parameters of the model, what each parameter means, and what estimates for each parameter were used (see main text and *Appendix III* for full details of how experiments were used to estimate these parameters).

The new model incorporates regulation by the CCO in two ways. First, Polo phosphorylates Cnn at multiple sites to promote scaffold assembly (22, 24, 26), so we make the simplifying assumption that Cnn molecules within the scaffold are stable when fully phosphorylated and must be fully dephosphorylated to be released into the cytoplasm. We allow a CCO regulated phosphatase activity to promote the dephosphorylation of the Cnn scaffold, while a CCO regulated kinase activity opposes this process by re-phosphorylating Cnn. Thus, the Cnn scaffold is stabilised in mitosis (when global CCO kinase activity is high and phosphatase activity is low) and destabilised at the end of mitosis and in early S-phase (when global CCO kinase activity is low and phosphatase activity is high). Second, Spd-2 appears to be phosphorylated by Cdk/Cyclins towards the end of each cycle, suppressing Spd-2’s ability to form a scaffold (41). We therefore allow rising CCO activity to suppress Spd-2 scaffold stability as the cycle progresses.

From these principles we derive two equations to describe how the amount of centrosomal Spd-2 and Cnn changes during each cycle of centrosome assembly (the model is summarised schematically in Fig. 4B; see *Appendix II* for full derivation):

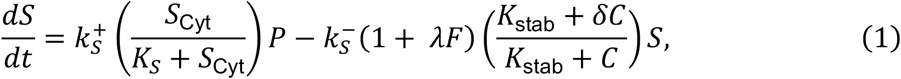

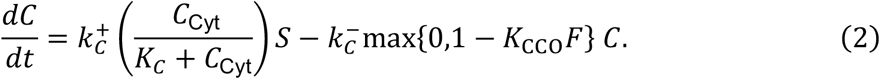

The first equation describes how the amount of Spd-2 scaffold, *S*, changes over time. The first term is the overall rate of scaffold generation, and the second term is the overall rate of scaffold loss through degradation or disassembly. Recruitment is assumed to be an enzymatic reaction, in which Polo at the centriole, *P*, binds to cytoplasmic Spd-2, *S*_Cyt_, in a reversible manner with dissociation constant *K*_*S*_. This Spd-2 is phosphorylated by the centriolar pulse of Polo to produce an unstable scaffolding form that is released from the enzyme complex at a rate *k*_*S*_^+^. The unstable Spd-2 scaffold has a disassembly rate, *k*_*S*_^−^, but it can be stabilised by binding to the Cnn scaffold, *C*, in a reversible manner with dissociation constant *K*_stab_. When bound to the Cnn scaffold, the Spd-2 scaffold has disassembly rate *k*_*S*_^−^*δ*, where *δ* < 1 is a parameter that measures the extent to which the Cnn scaffold stabilises the Spd-2 scaffold. The rate of disassembly of the Spd-2 scaffold is influenced by the CCO, which is expressed here as the ratio of cell cycle kinase/phosphatase activity in the system, *F*. The sensitivity of the Spd-2 OFF rate to the cell cycle kinase/phosphatase ratio is measured by the parameter *λ*.

The second equation describes how the amount of Cnn scaffold changes over time. The first term is the overall rate of scaffold generation, and the second term is the overall rate of scaffold loss through degradation or disassembly. Cnn is assumed to be recruited enzymatically by binding from the cytoplasm, *C*_Cyt_, to the Spd-2 scaffold in a reversible manner before being released in scaffold form. The dissociation constant in this enzymatic recruitment process is *K*_*C*_, and the rate of release is *k*_*S*_^+^. The off rate of Cnn is given by *k*_*S*_^−^, but we allow this to be influenced by the cell cycle kinase/phosphatase ratio according to the expression max{0,1 − *K*_CCO_*F*}. This is due to the competition between CCO kinase activity phosphorylating and stabilising the scaffold, and CCO phosphatase activity dephosphorylating and destabilising the scaffold. Here, *K*_CCO_is the ratio of the rate of phosphorylation to the rate of dephosphorylation—a constant parameter that measures the reaction rates of these two processes—while *F* is a function that measures the varying ratio of kinase to phosphatase activity as the cell cycle progresses.

### Generating the model inputs

Our model contains two inputs: (1) the initial pulse of centriolar Polo activity, *P*, and (2) the activity of the CCO, which we express here as the ratio of cell cycle kinase/phosphatase activity, *F*. We previously characterised the centriolar Polo pulse and modelled its generation by the centriole protein Ana1/CEP295, which is a major recruiter of Polo to centrioles (48). Here, for simplicity, we generated an offset sinusoidal pulse function to approximate this activity. To generate *F*, we used a FRET-based Cdk1 activity sensor that has previously been validated in early *Drosophila* embryos (49). This cytoplasmic sensor is phosphorylated by Cdk1/Cyclins and is dephosphorylated by opposing phosphatases, so its emission ratio provides a broad readout of the balance between global CDK kinase and opposing phosphatase activity as embryos progress through the nuclear cycle (Fig. S2).

### Experimental estimation of model parameters

Our model contains 10 parameters (Equations 1 and 2) that we proceeded to estimate experimentally (Fig. 4C) (see *Appendix III*). To estimate the dissociation constants *K*_*S*_and *K*_*C*_, we measured how the centrosomal recruitment of Spd-2 and Cnn changes as their cytoplasmic concentration changes. We used a series of CRISPR knock-ins and heterologous transgenic lines to express NG- or GFP-tagged Spd-2 or Cnn at different levels in embryos (Fig. S3). We used fluorescence correlation spectroscopy (FCS) to measure the cytoplasmic concentration of the tagged protein in each line at the start of NC12 (Fig. 5A), and confocal microscopy to measure its centrosomal recruitment dynamics (Fig. 5B). As the cytoplasmic concentration increased, both the rate at which each protein accumulated at centrosomes and the amount recruited increased. In the model, the recruitment rate of each protein depends on its cytoplasmic concentration through a Michaelis–Menten term, *C*/(*K* + *C*). Fitting this term to the relationship between cytoplasmic concentration (Fig. 5C) gave estimates of *K*_*S*_ = 270nM and *K*_*C*_ = 70nM (Fig. 4C).

**Fig. 5.**
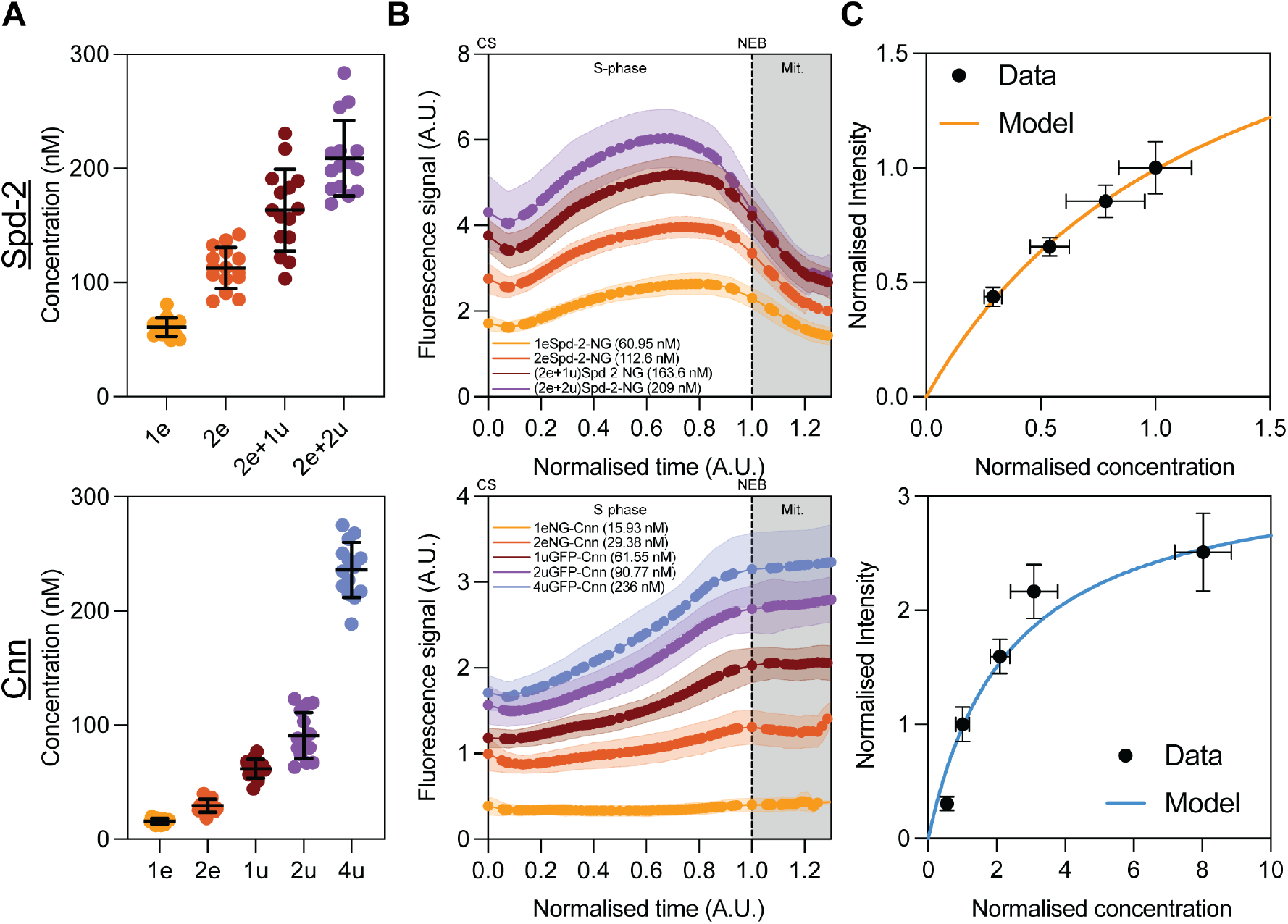
The centrosomal recruitment of Spd-2 and Cnn scales with their cytoplasmic concentration. **(A)** Cytoplasmic concentrations of Spd-2-NG (top) and either NG-Cnn or GFP-Cnn (bottom), measured by FCS at the start of NC12 in embryos expressing different amounts of each fusion protein in a mutant background that lacks untagged endogenous protein. “e” indicates an endogenous CRISPR knock-in allele and “u” a ubiquitin promoter-driven transgene; the number gives the copy number. The 1e and 2e Cnn lines express NG-Cnn, and the 1u, 2u and 4u lines express GFP-Cnn. Each point is one embryo. Bars show mean ± SD. **(B)** Mean centrosomal fluorescence of Spd-2-NG (top) and Cnn (bottom) during NC12 in the lines shown in (A). The mean cytoplasmic concentration of each line is given in the key. Time is normalised (0 = centrosome separation, 1 = NEB; grey shading, mitosis). Shaded bands show SD. n >1000 centrosomes from N = 12-15 embryos per line. NG and GFP differ in brightness, so the NG-Cnn traces were rescaled onto the GFP-Cnn scale using a calibration between 1u NG-Cnn and 1u GFP-Cnn lines with matched cytoplasmic concentrations (Fig. S3B,C). **(C)** Normalised peak centrosomal intensity plotted against normalised cytoplasmic concentration for Spd-2 (top) and Cnn (bottom). Both axes are normalised so that the line closest to endogenous expression levels, judged by Western blot (Fig. S3A), equals 1. Error bars show SD. Lines are fits of the Michaelis–Menten recruitment term in Equations 1 and 2, giving *KS*= 270nM and *KC*= 70nM.

We next used the data from our earlier FRAP measurements (Fig. 2) to constrain the Spd-2 and Cnn scaffold loss terms. We used the early-cycle Spd-2 OFF rate to estimate basal Spd-2 loss, *k*_*S*_^−^, since Cnn stabilisation and CCO destabilisation are low in this time. During the mid to late cycle, we assume that enough Cnn scaffold has been generated that most of the Spd-2 scaffold being generated by the centrioles becomes stabilised, so any variation in Spd-2 stability is solely due to cell cycle effects. Therefore, we use this variation to estimate the magnitude of the CCO’s influence on Spd-2 scaffold stability, *λ*. We use the late-cycle OFF rate to estimate the stabilising effect of Cnn, *δ*, since the Spd-2 is fully stabilised by Cnn at this time, and we estimate the dissociation between the Spd-2 and Cnn scaffolds, *K*_stab_, using the time it takes for the OFF rate to transition between the unsaturated regime in the early cycle and the Cnn-saturated regime in the mid-to-late-cycle. The early cycle OFF rate of Cnn describes the basal Cnn loss, *k*_*C*_^−^, since the stabilising influence of the CCO on the Cnn scaffold is minimal at this time, and the rate at which the Cnn OFF rate approaches zero informs the magnitude of the CCO’s influence on the Cnn scaffold stability, *K*_CCO_. The physical values of the remaining incorporation parameters, *k*_*S*_^+^ and *k*_*C*_^+^, cannot be measured since this would require measuring the number of Spd-2 and Cnn molecules in the centrosome, which we were not able to do. However, for model consistency, we choose these parameters such that Spd-2 and Cnn reach their normalised peak values during nuclear cycle 12. Full parameter derivations and dimensional conversions are given in *Appendix III*.

With these parameter constraints, the model generally reproduces the major behaviours of the Spd-2 and Cnn scaffolds well. A single Polo pulse generates a transient Spd-2 recruitment response, while Cnn continues to accumulate towards a plateau even after Spd-2 levels have begun to decline (Fig. 6A) (41). Analysis of the modelled Spd-2 and Cnn OFF rates show that the model broadly captures the trend of how these parameters change during the cycle (Fig. 6B). The model reproduces the early-to-mid-cycle decrease in the Spd-2 OFF rate, which in the model is due to the stabilising effect of the newly forming Cnn scaffold, as well as the later mid-cycle-to-mitosis rise in the Spd-2 OFF rate, which in the model is due to the rising CCO activity destabilising the Spd-2 scaffold. The dramatic fall in the Cnn OFF rate in mid-S-phase is also captured well, which in the model is due to the rising CCO activity inactivating the Cnn-scaffold destabilising phosphatases.

**Fig. 6.**
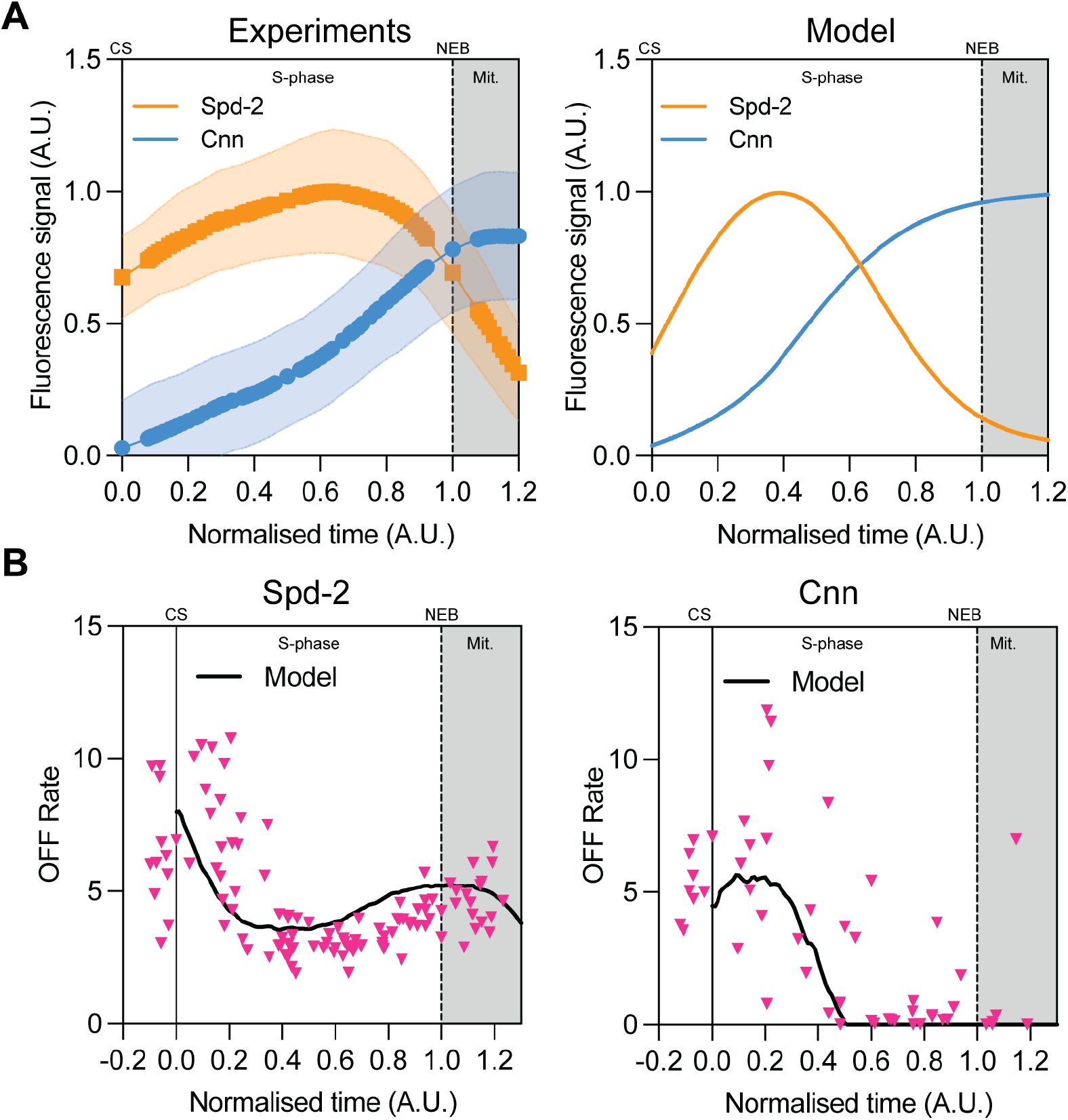
The model reproduces well the main features of Spd-2 and Cnn scaffold assembly. **(A)** Left graph (experiments) show how centrosomal Spd-2 or Cnn levels change over time during NC12. Right graph (Model) shows how the model predicts Spd-2 and Cnn levels change over time during a nuclear cycle. The data in the left graph were replotted from Fig. 1C from Wong et al., (21), with time renormalised (0 = centrosome separation, 1 = NEB; grey shading, mitosis). Shaded bands show SD. The model was driven by a single sinusoidal pulse of centriolar Polo activity, with CCO activity (F) taken from the Cdk1 FRET sensor data (Fig. S2). Model parameters are listed in Fig. 4C. **(B)** Spd-2 (left) and Cnn (right) OFF rates from the model (black lines), overlaid on the FRAP-derived OFF rates from Fig. 2C (magenta triangles). The parameters that set the scaffold loss terms were estimated from these data (Fig. 4C). CS, centrosome separation; NEB, nuclear envelope breakdown.

### Cell-cycle state controls Spd-2 and Cnn scaffold dynamics

In our model, the changing dynamics of the Spd-2 and Cnn scaffolds during the nuclear cycle are largely driven by changes in CCO activity. We therefore wanted to directly test the role of the CCO in regulating scaffold dynamics, and to ask whether our model could accurately predict how perturbations to the CCO would affect scaffold assembly. We first injected double-stranded RNAs targeting the three embryonic mitotic cyclins, Cyclin A, Cyclin B and Cyclin B3, into embryos expressing either Spd-2-GFP or NG-Cnn. This treatment arrests embryos in a low CCO activity state prior to S-phase entry (50). The centrosomes in these embryos no longer generated pulses of Polo recruitment (*green line*, Fig. 7A), supporting our hypothesis that CCO activity is required to initiate these pulses (18). Instead, centrosomal Polo levels gradually declined to an intermediate level.

**Fig. 7.**
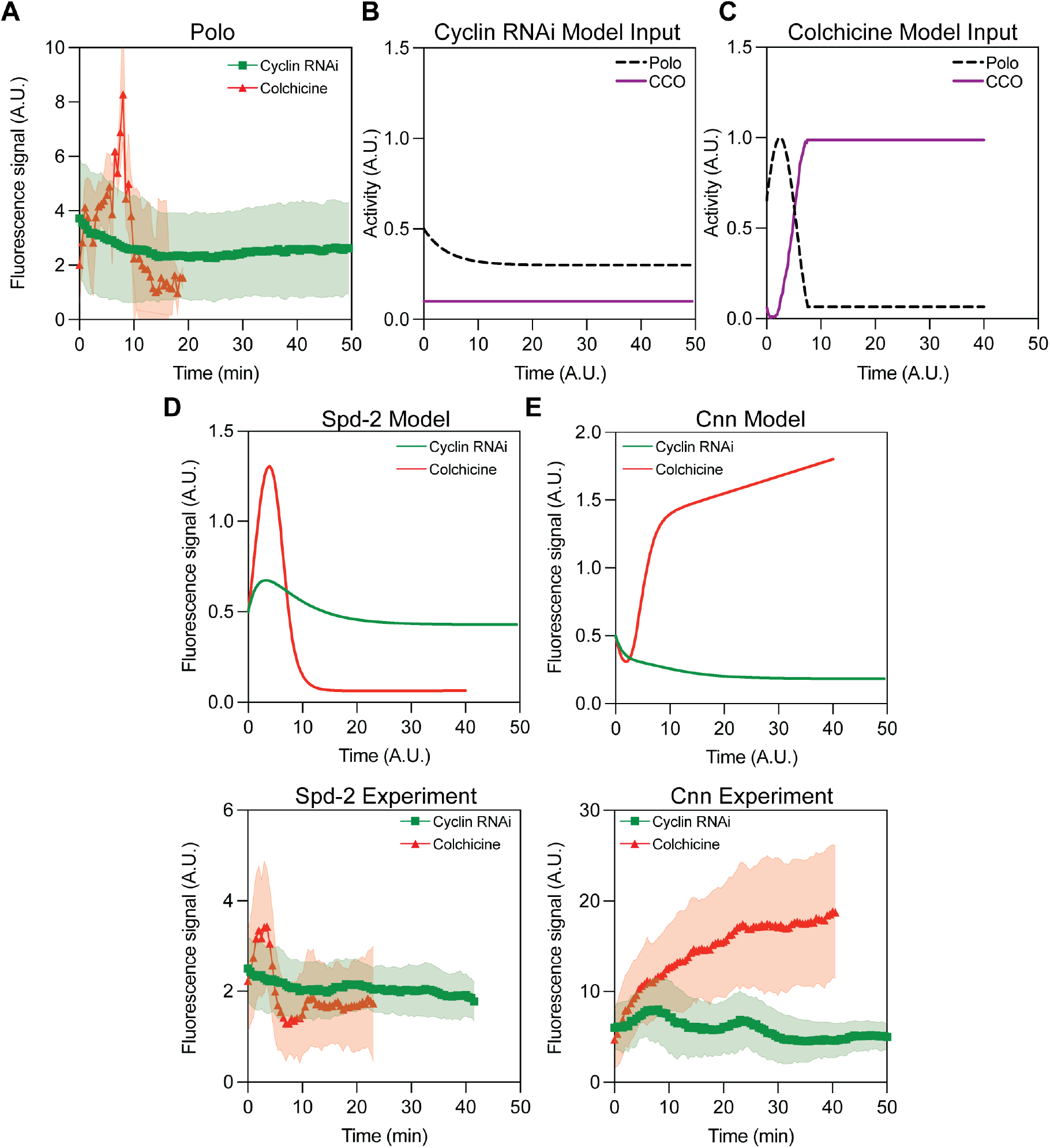
The model reproduces well how the Spd-2 and Cnn scaffolds respond to CCO perturbations. **(A)** Graph shows how centrosomal Polo levels change over time in embryos arrested with low CCO activity (by Cyclin RNAi, green line) or with high CCO activity (by colchicine injection, red line). Note that in embryos arrested in mitosis, Polo rapidly accumulates on the kinetochores and is largely lost from centrosomes; this makes it impossible to accurately track (and so measure) centrosomes during the prolonged mitotic arrest. Nevertheless, qualitatively, centrosomal levels of Polo remain low throughout the arrest period. **(B)** Graphs show the model inputs for the centriole Polo activity pulse (black dotted lines) and global CCO activity (purple lines) that we used to model embryos arrested with low CCO activity (Cyclin RNAi, left graph) or with high CCO activity (Colchicine, right graph). **(D,E)** Graphs show the model prediction (top graphs) or experimental data (bottom graphs) of how the Spd-2 (left) or Cnn (right) scaffolds behave in embryos arrested with low CCO activity (green lines) or high CCO activity (red lines).

We used this information to adjust the Polo and CCO inputs to our model, allowing centrosomal Polo activity to decay gradually to a steady intermediate level, and holding CCO activity at a constant low-level (Fig. 7B). Under these conditions, the model predicts that the Spd-2 and Cnn scaffolds should both gradually decline to a low steady state level at which their ON and OFF rates are balanced (*green lines*, Model graphs, Fig. 7D,E). Both predictions largely agreed with our experimental observations of Spd-2 and Cnn scaffold behaviour in Cyclin RNAi embryos (*green lines*, Experiment graphs, Fig. 7D,E).

We next asked what happens to the Spd-2 and Cnn scaffolds if CCO activity is held high. To do this, we injected embryos with colchicine at the start of S-phase, which disrupts subsequent spindle assembly, activates the mitotic checkpoint, and arrests embryos in a mitotic state. After injection, we monitored embryos as they proceeded through one round of relatively normal Spd-2 and Cnn scaffold assembly before arresting in the next mitosis. In these embryos, Polo levels underwent one relatively normal round of recruitment in the run up to mitosis and then dropped dramatically as the embryo arrested in mitosis (*red line*, Fig. 7A).

We again used this information to adjust the Polo and CCO inputs to our model, allowing centrosomal Polo activity to undergo a normal pulse of recruitment followed by a rapid decay to a low steady-state level, and CCO activity to rise normally during the early cycle and to then be maintained at a constant high-level (Fig. 7C). Under these conditions, the model predicts that the Spd-2 and Cnn scaffolds exhibit very different behaviours. The Spd-2 scaffold initially assembles but then disassembles to a low steady state level (*red line*, Spd-2 Model graph, Fig. 7D), whereas the Cnn scaffold assembles rapidly during the normal S-phase but then continues to grow more slowly during the prolonged mitotic arrest (*red line*, Cnn Model graph, Fig. 7E). Again, both predictions largely agreed with our experimental observations of Spd-2 and Cnn scaffold behaviour in these embryos (*red lines*, Experiment graphs, Fig. 7D,E).

Together, these experiments indicate that the CCO differentially regulates the dynamics of the two scaffolds—high CCO activity destabilises the Spd-2 scaffold but stabilises the Cnn scaffold— and that our model captures these behaviours reasonably well.

### The CCO and Cnn modulate the stability of the Spd-2 scaffold

We next wanted to test whether our model faithfully reproduced two key aspects of Spd-2 scaffold behaviour: How the Cnn scaffold stabilises the Spd-2 scaffold, and how rising CCO activity destabilises the Spd-2 scaffold. These effects can be analysed experimentally by halving the genetic dose of Cnn (Experiment, Fig. 8A) or by examining the behaviour of a Spd-2 mutant that cannot be phosphorylated by Cdk/Cyclins (Spd-2-Cdk20A) (41) (Experiment, Fig. 8B). These effects can be simulated in the model by halving the amount of Cnn in the system (Model, Fig. 8A) or by setting the dependence of the Spd-2 OFF rate on the cell cycle kinase/phosphatase ratio (*λ*—Equation 1) to zero (Model, Fig. 8B). The model captures these experimental behaviours well. Note that in these graphs, to more fairly compare model and experiment, we subtracted the minimum value from each centrosomal trace and normalised the resulting curves to the WT peak. This focuses the data on new material incorporated during the cycle rather than on any residual centrosomal scaffold inherited from earlier cycles—which would not be captured in our model where growth is assumed to start from zero.

**Fig. 8.**
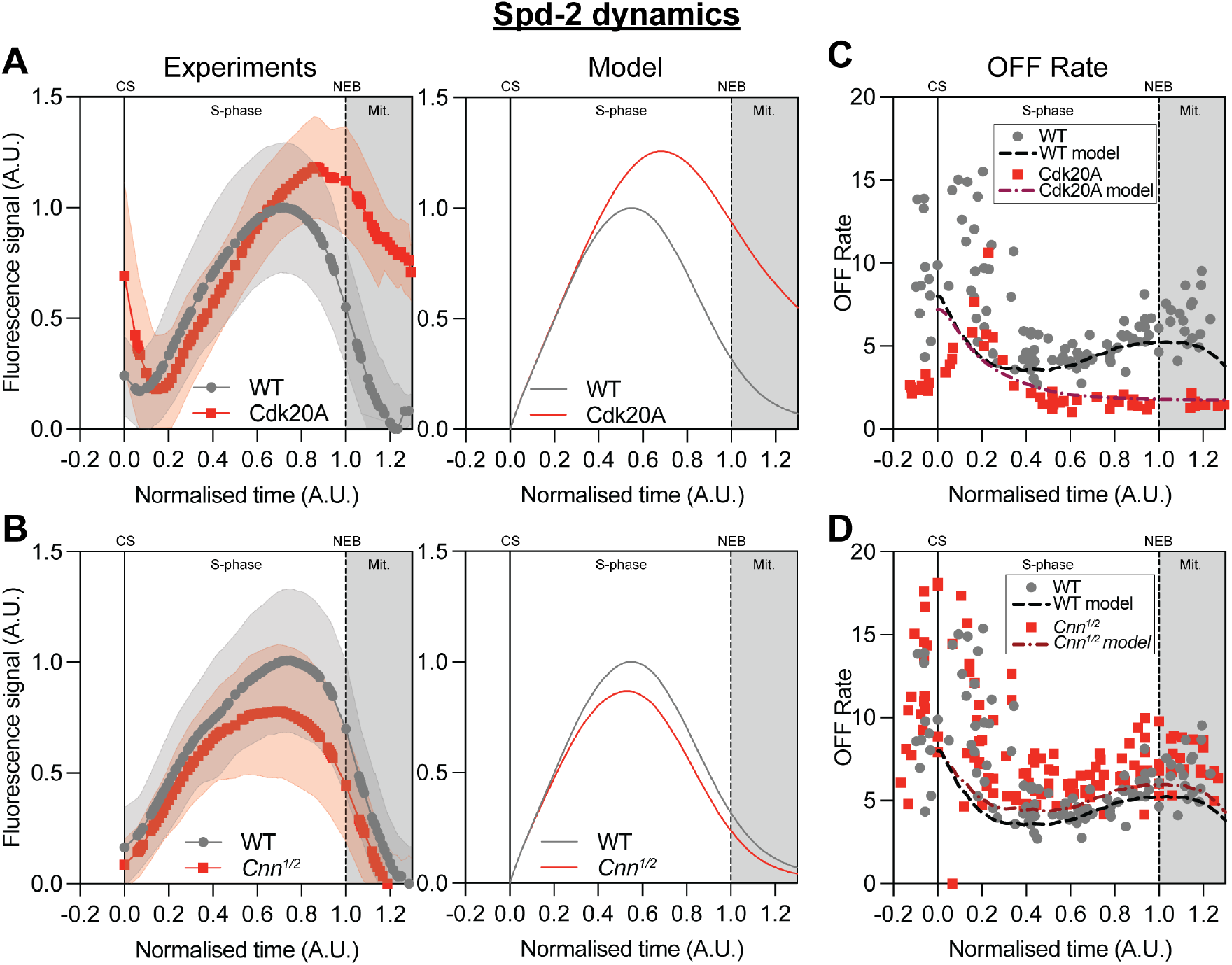
The model reproduces well how the Spd-2 scaffold responds to perturbations in its sensitivity to Cdk/Cyclin phosphorylation and to Cnn levels. **(A,B)** Left graphs (Experiments) show how centrosomal Spd-2 levels change over time in embryos expressing a form of Spd-2 that cannot by phosphorylated by Cdk/Cyclins (Cdk20A) (A) or in which the genetic dose of Cnn has been halved (B). Right graphs (Model) show model predictions of how the Spd-2 scaffold responds to these changes. **(C,D)** Graphs show how the Spd-2 OFF rates change during NC12 (each individual data point is calculated from a FRAP experiment, as shown in Fig. 2) for WT Spd-2 or Spd-2-Cdk20A (top graph) or for WT Spd-2 in WT or *Cnn* half-dose (*Cnn^1/2^*) embryos (bottom graph). The overlayed dotted lines show how the model predicts these perturbations should change Spd-2 OFF rates.

The model predicts that the main reason halving the dose of Cnn reduces centrosomal Spd-2 levels is because this increases the Spd-2 OFF rate (*dotted lines*, Fig. 8C), and a FRAP analysis confirmed that this matched the experimental OFF rate data well (*data points*, Fig. 8C). In contrast, the model predicts that the main reason that Spd-2-Cdk20A accumulates at centrosomes towards the end of the cycle is that the Spd-2-Cdk20A OFF rate no longer increases as the embryo prepares to enter mitosis (*dotted lines*, Fig. 8D), and a FRAP analysis confirmed that this modelled the data well (*data points*, Fig. 8D). These analyses indicate that our model captures well how Cnn levels and the CCO influence the behaviour of the Spd-2 scaffold.

### How Spd-2 and Polo modulate the stability of the Cnn scaffold

We next wanted to test whether our model faithfully reproduced how the Cnn scaffold is influenced by the levels of Spd-2 and Polo in the embryo. As intuitively expected, experimentally halving the genetic dose of Spd-2 led to a decrease in the amount of Cnn recruited to centrosomes (Experiment, Fig. 9A). Note that for this analysis, as above, we normalised the experimental data to focus on new material incorporated during the cycle. This behaviour was captured reasonably well by halving the amount of Spd-2 in our model (Model, Fig. 9A).

**Fig. 9.**
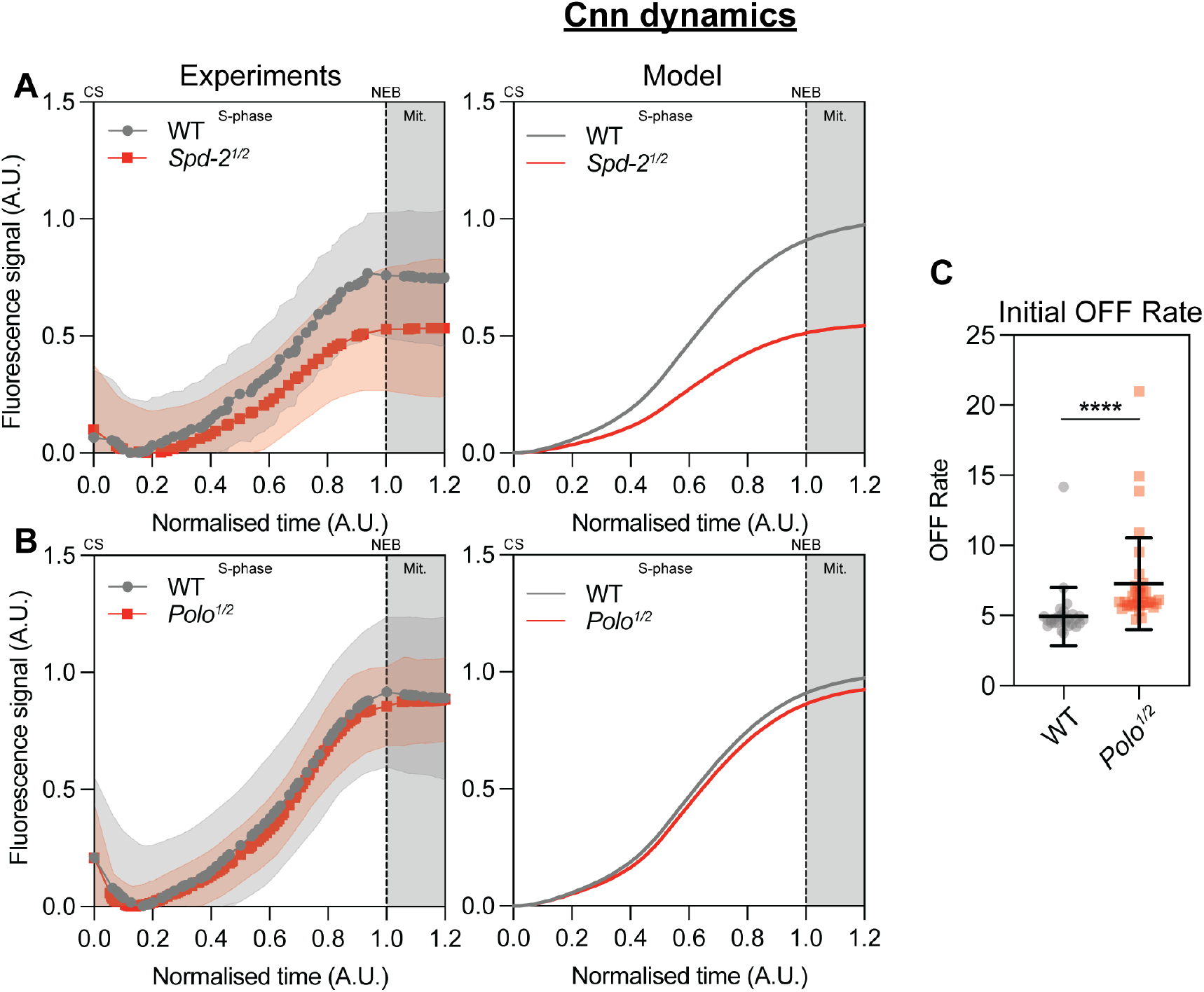
The model reproduces well how the Cnn scaffold responds to lowering the dose of Spd-2 and Polo. **(A,B)** Left graphs (Experiments) show how centrosomal Cnn levels change over time in WT embryos or in embryos in which the genetic dose of *Spd-2* (top graph) or *Polo* (bottom graph) has been halved (*Spd-2^1/2^* and *Polo^1/2^*, respectively). Right graphs (Model) show model predictions of how the Cnn scaffold responds to these changes. (C) Graph shows the Cnn OFF rate early in NC12 in WT embryos and in embryos in which the genetic dose of *Polo* has been halved.

To our surprise, we found that experimentally halving the genetic dose of Polo did not appreciably perturb the amount of new Cnn material that added into centrosomes during the nuclear cycle (Experiment, Fig. 9B; data normalised as described above). Interestingly, however, a FRAP analysis revealed that halving the genetic dosage of Polo did increase the initial Cnn OFF rate early in the cycle by ∼33%, as expected because rising levels of Polo activity in the cytoplasm are required to counter the high levels of phosphatase activity that destabilises the Cnn scaffold at the end of mitosis (Fig. 9C).

Modelling the effect of decreasing the dose of Polo in our model is not straightforward, as this perturbation potentially influences several elements (e.g., the centriolar Polo pulse input, CCO activity, the stability of the Cnn scaffold). We found, however, that our model could reproduce this surprising behaviour (Model, Fig. 9B) if we allowed halving the amount of Polo to reduce the rate of phosphorylation, and therefore the stability, of the Cnn scaffold by 33%—as indicated experimentally by the increase in Cnn OFF (Fig., 9C)—but allowed the input pulse of Polo recruitment to the centriole to remain unaltered. This last point seems plausible, as the levels of PLK1 activity required to initiate spindle assembly is low compared to the levels required for events that occur later in mitosis (51). On inspection of the model, we can see that this behaviour occurs because the OFF rate of Cnn is near to zero for much of the cycle. Thus, the effect of reducing Polo on Cnn loss primarily affects the disassembly phase between cycles, and so the amount of new material incorporated throughout the cycle is minimally affected by Polo concentration.

## Discussion

The dramatic expansion of the PCM prior to mitosis seems to be a universal feature of centrosomes in dividing cells. In most species, centrosomes start growing sometime before mitosis, largely stop growing during mitosis, and then disassemble at the end of mitosis (13, 14, 41, 52). It is widely recognised that the cell cycle oscillator (CCO) triggers mitotic centrosome growth prior to mitosis and centrosome disassembly at the end of mitosis, but it is less clear why centrosomes stop growing during mitosis. The prevailing model, largely based on experiments in the worm embryo, has been that centrosomes stop growing because they run out of a limiting component (29, 42). But recent studies in *Drosophila* embryos suggested an alternative possibility: that high CCO activity slows centrosome growth during mitosis. Here, we provide strong evidence that this is the case.

Our experiments and modelling suggest that the CCO influences mitotic centrosome growth in fly embryos by differentially regulating the stability of two key scaffolds that normally support mitotic centrosome growth: Spd-2/CEP192 and Cnn/CDK5RAP2. Early in the cycle, centrioles start to assemble dynamic Spd-2 and Cnn scaffolds. But, as the cycle progresses and CCO activity rises, the Spd-2 scaffold is destabilised, in part by the Cdk/Cyclin-dependent phosphorylation of Spd-2. This slows Cnn scaffold growth (which is dependent on the Spd-2 scaffold), but the existing Cnn scaffold is stabilised by the CCO-dependent inactivation of cell cycle phosphatases—locking Cnn in a low-dynamic state that is independent of Spd-2. Thus, centrosomes dramatically slow their growth as the embryos enter mitosis, but the existing Cnn scaffold is maintained.

Given that flies and worms use such a similar set of proteins to build their mitotic PCM it is perhaps surprising that they appear to use such different mechanisms to limit centrosome growth (CCO regulation in flies; a limiting component in worms). We believe, however, that our model can at least qualitatively explain the kinetics of centrosome growth in the early worm embryo. Recent experiments have shown that SPD-2 and Polo/PLK-1 are required for the growth of the SPD-5 scaffold in worm embryos but, once assembled, the SPD-5 scaffold is no longer dependent on SPD-2 (32). Moreover, there is strong evidence that the SPD-5 scaffold is initially dynamic but gradually transitions to a more stable, low-turnover state in a phosphorylation dependent manner (30, 31, 34, 53). Moreover, at the end of mitosis the rise in phosphatase activity leads to the rapid disassembly of the SPD-5 scaffold (15, 16).

These observations in worms fit very well with our model of scaffold assembly in flies, and we suggest that there may be no need to invoke a limiting component to explain centrosome growth kinetics in the worm embryo. A recent analysis suggested that a limiting component may not explain centrosome size regulation in the worm embryo system (54). It will be interesting to determine whether CEP192 and CDK5RAP2/CEP215, the human homologues of Spd-2/SPD-2 and Cnn/SPD-5, respectively, exhibit similar dynamic behaviours. A recent FRAP analysis concluded that CDK5RAP2/CEP215 was immobile in interphase, but mobile in mitosis, although this analysis did not consider the possibility that centrosome growth might account for some of the fluorescence recovery observed during mitosis (55).

We stress that the differential regulation of the Spd-2 and Cnn scaffolds is certainly not the only way that the CCO regulates centrosome growth. As discussed above, it is well established that centrosome maturation initiates at a specific point in the cell cycle (12), and in fly embryos this seems to occur when centrioles generate a pulse of Polo activity (21). In support of this latter possibility, we found here that centrosomes no longer generate such Polo pulses if embryos are held in a low CCO activity state by the depletion of Cyclins A, B and B3. Moreover the rapid disassembly of the PCM at the end of mitosis is also clearly a cell cycle regulated event, which in worm embryos is triggered by the activation of PP2A at the end of mitosis (15, 16). The relevant phosphatase(s) in fly embryos has/have yet to be identified.

Although *C. elegans* and *Drosophila* are thought to be separated by ∼800-1000 million years of evolution (56), it is intriguing that they both adopt a “two-scaffold” solution to grow their mitotic centrosomes. Why not just use one scaffold? We speculate that this may be because centrosome growth is a demanding process. The tiny centrioles must support the rapid growth of the much larger PCM, while at the same time the two centrosomes in a somatic cell (or the many hundreds of centrosomes in a syncytial *Drosophila* embryo) must grow to the same size. Mathematical modelling has shown that it is very difficult to satisfy both requirements at the same time, as the autocatalytic mechanisms required to generate rapid growth are generally incompatible with robust size regulation (2, 43). Having two scaffolds with different dynamic properties may solve this problem. In fly embryos, the growth of the Cnn scaffold is catalytic not because Cnn can recruit itself but because the Cnn scaffold stabilises the Spd-2 scaffold from which it is generated. Thus, the growth of the Cnn scaffold remains limited by the size of the Spd-2 scaffold, which in turn is limited by the size of the centriole.

This two-scaffold solution allows for rapid centrosome growth, as the expansion of the Spd-2 scaffold around the mother centriole provides a greater surface area to generate more Cnn scaffold as centrosomes grow. But it also allows for robust size regulation, as growth is ultimately regulated by the rate at which centrioles can generate the Spd-2 scaffold, rather than by the amount of Cnn scaffold present at each centrosome—as would be the case if the Cnn scaffold was genuinely autocatalytic. In such a system it can be the size of the mother centriole that ultimately determines the relative size of any centrosomes. This may be why it is so important that centrioles grow to a consistent size. We are currently developing mathematical models to formally test this hypothesis.

## Materials and Methods

### *Drosophila* stocks and husbandry

All Drosophila melanogaster stocks used in this study are listed in Table S1. Flies were maintained at 18°C or 25°C in plastic vials or bottles on standard cornmeal medium (0.68% agar, 2.5% yeast, 6.25% cornmeal, 3.75% molasses, 0.42% propionic acid, 0.7% ethanol and 0.14% Tegosept). For embryo collections, adult flies were transferred to cages containing fruit-juice agar plates (40% cranberry–raspberry juice, 2% sucrose and 1.8% agar) supplemented with a drop of yeast paste. Fly husbandry followed procedures designed to minimize plastic waste (57).

### Embryo collection and mounting

Embryos were collected from fruit-juice agar plates after a 60-min egg lay at 25°C, unless stated otherwise. Embryos were dechorionated by hand and mounted on a strip of glue in 35-mm glass-bottom dishes with 14-mm microwells (P35G-1.5-14-C, MatTek). They were then desiccated for 1–5 min at 25°C and covered with Voltalef H10S PCTFE oil (Arkema) to prevent further desiccation.

### dsRNA synthesis and injection

dsRNAs against Cyclin A, Cyclin B and Cyclin B3 were synthesized essentially as described previously (50). dsRNA was transcribed with the MEGAscript T7 Transcription Kit (AM1334, Invitrogen) and purified with an RNA Clean & Concentrator kit (Zymo Research). Complementary strands were annealed at 67.5°C for 30 min and then cooled slowly to room temperature over 60 min. Purified dsRNA was stored at −70°C. For injection, embryos were collected after a 30-min egg lay, and a mixture of all three dsRNAs was injected at a final concentration of 200 ng/ μl each. Embryos were incubated for a further 30 min at 25°C before imaging.

### Colchicine injection

Embryos were desiccated for 5–10 min, depending on ambient humidity, and covered with Voltalef oil. They were then injected with 100 μg/ml colchicine, diluted in Schneider’s medium from a 1 mg/ml stock in dimethyl sulfoxide (DMSO).

### Immunoblotting

Embryos were fixed in methanol and stored at 4°C for at least one night before being rehydrated with three 15-min washes in PBT (PBS + 0.1% Triton X-100) and processed for SDS–PAGE and immunoblotting as described previously (25). We used the following primary antibodies: rabbit anti-Cnn (1:2000) (58), rabbit anti-Spd-2 (1:1000) (59), rabbit anti-GAGA factor (1:500) (60) and mouse anti-actin (A3853, Sigma-Aldrich; 1:500). Secondary antibodies were horseradish peroxidase– conjugated donkey anti-rabbit (NA934V) and sheep anti-mouse (NA931) IgG (both Cytiva, supplied by VWR International), used at 1:3000–1:5000. Signal was detected with diluted SuperSignal ECL substrates (Thermo Fisher Scientific).

### Spinning-disk confocal microscopy

Living embryos were imaged at 23°C on a PerkinElmer ERS spinning-disk confocal system mounted on a Zeiss Axiovert 200M microscope and controlled by Volocity software (PerkinElmer). All images were acquired with a 63×/1.4 NA oil-immersion objective, using Immersol 518 F oil (Carl Zeiss; refractive index 1.518) to minimize spherical aberration. Images were captured on an Orca ER CCD camera (Hamamatsu Photonics) at a gain of 200. The system was equipped with 405-, 488-, 561- and 642-nm solid-state lasers (Oxxius). Two-colour samples were imaged with the UltraVIEW ERS “Emission Discrimination” setting, using a 520-nm long-pass filter for the green channel and a 620-nm long-pass filter for the red channel. For two-colour imaging, the red channel was acquired before the green channel at each z-slice. Z-stacks were acquired at 0.5-μm intervals. The number of slices, time step, laser power and exposure varied by experiment.

For FRAP experiments, circular regions of interest 3.5–4 μm in diameter were drawn around individual centrosomes. These were bleached with 20 iterations of the 488- and 561-nm lasers at 50% power. In some embryos, different centrosomes were bleached at different times.

### Fluorescence correlation spectroscopy (FCS)

FCS was performed essentially as described previously (61, 62). Measurements were made on a Zeiss LSM 880 confocal microscope controlled by Zen Black software. We used 488-nm argon laser excitation, a GaAsP photon-counting detector (491–544 nm) and a C-Apochromat 40×/1.2 W objective with the pinhole set to 1 Airy unit. At the start of each experimental day, the correction collar was adjusted to maximize the counts per molecule (CPM) of a fluorescent dye solution. The effective observation volume (V_eff_) had been estimated previously at ∼0.25 fl, using Alexa Fluor 488 diffusion measurements and subresolution fluorescent beads. No photobleaching was detected under these conditions. The stage was held at 25.0–26.0°C with a Zeiss XL incubator.

For each measurement, six consecutive 10-s recordings were taken in the cytoplasm at the centrosomal plane. Recordings were discarded if a centrosome or yolk granule entered the observation volume. Measurements were kept only if at least four of the six recordings were valid. For time courses spanning whole nuclear cycles, an embryo was kept only if every timepoint passed these criteria.

Autocorrelation curves were analysed in FoCuS-point (63). Curves were fitted with models containing one or two diffusing species and different fluorophore dark states. The structural parameter (AR = ω_z_/ω_xy_) was fixed at 5. The anomalous diffusion parameter (α) was chosen separately for each protein using the Bayesian information criterion (64). After background correction, we calculated cytoplasmic concentration, diffusion coefficient and CPM. Outliers were removed with the ROUT method in GraphPad Prism (Q = 1%).

### Cdk1 FRET imaging

To measure Cdk1 activity, we imaged embryos expressing the Cdk1-FRET sensor (49) and Jupiter-mCherry on a Zeiss LSM 880 confocal microscope fitted with a spectral GaAsP detector and controlled by ZEN Black v2.3. Images were acquired with a 20×/0.8 NA Plan-Apochromat air objective. CFP was excited at 458 nm with a multiline argon laser, and CFP (471–508 nm) and YFP (543–606 nm) emissions were recorded at the same time. Jupiter-mCherry was excited with a 561-nm solid-state laser and detected at 615–735 nm. The pinhole was set to 1.13 Airy units. Images were 512 × 512 pixels, with a voxel size of 0.415 × 0.415 × 0.8 μm. Stacks of five z-slices were acquired every 8 s, at a scan time of 1.26 s per slice.

### Image analysis

Centrosome detection and intensity measurements. Unless stated otherwise, raw z-stacks were maximum-intensity projected, and uneven background illumination was corrected using a custom Python script (21). Centrosomes were detected and tracked over time using either TrackMate (65) in Fiji or in a Python pipeline. In the Python pipeline, centrosomes were detected using the Crocker–Grier centroid-finding algorithm (66) as implemented in TrackPy (67). Signal-to-background thresholds were set using Otsu’s method (68). Centrosomes were then linked between frames using LapTrack (69). Centrosome fluorescence intensity was measured as the total intensity after subtraction of local background.

#### FRAP analysis

Before projection and background correction, time series were corrected for photobleaching by fitting an exponential decay function. Bleached centrosomes were then tracked manually using a custom annotation tool, adapted from a software package developed for mobile robotics (70). A custom Python script extracted fluorescence intensities at each tracked centrosome over time in each embryo, as described previously (21). The models fitted to these curves, their derivations and the parameter-estimation procedures are described in *Appendix I*.

#### Cdk1 FRET analysis

Cdk1 activity was measured as the FRET ratio, as described previously (49). z-stacks were maximum-intensity projected, leaving out slices with high autofluorescence from the embryo edge. The FRET ratio was calculated as the mean YFP signal divided by the mean CFP signal within a region drawn inside the embryo. The FRET ratio drifted downwards over time, and the same drift was seen with a control sensor in which the phosphosite was mutated to alanine. To correct for this drift, we fitted an exponential decay through the trough of each oscillation in GraphPad Prism (v11) to define a baseline, then subtracted it from the raw FRET ratio. Cell-cycle timing was defined using the Jupiter-mCherry signal. For each embryo, nuclear cycle 12 was time-normalised, with disassembly of the telophase spindle remnant in cycle 11 taken as the start and the same event in cycle 12 taken as the end. FRET ratios at fixed normalised time points were interpolated from the measured data using a custom Python script, so that data from different embryos could be averaged.

### Statistical analysis

Statistical tests, sample sizes and definitions of centre and dispersion are given in the relevant figure legends. All statistical analyses were performed in GraphPad Prism.

## Data and code availability

All raw image data will be deposited in the BioImage Archive (71) on publication. Custom analysis scripts will be made available on GitHub on publication. Existing analysis code was updated to current software versions with the assistance of Codex ChatGPT.

## Acknowledgements

We thank Stefano De Talia for the CDK FRET sensor and members of the Raff Laboratory for many helpful discussions and comments. We thank the Light Microscopy Facility, part of the Dunn School Bioimaging Facility, managed by Alan Wainman, for providing support with live-cell imaging. We also thank Saroj Saurya for ensuring that our laboratory work and reagent use adhered to strict environmental sustainability standards.

## Author contributions

Conceptualization: SSW, ZMW, JWR

Methodology: SSW, ZMW, ELR, KYC, TLS, JWR

Investigation: SSW, ZMW, ELR, JWR

Resources: SSW, ZMW, ELR, KYC, TLS, JWR

Data curation: SSW, ZMW, ELR, JWR

Formal analysis: SSW, ZMW, JWR

Software: SSW, KYC

Validation: SSW, ZMW, ELR, JWR

Visualization: SSW, ZMW, JWR

Project administration: SSW, ZMW, JWR

Supervision: SSW, ZMW, JWR

Writing – original draft: SSW, ZMW, JWR

Writing – review & editing: SSW, ZMW, ELR, KYC, TLS, JWR

## Data and materials availability

All data needed to evaluate the conclusions in the paper are present in the paper and/or the Supplementary Materials. All raw data will be made available in an appropriate public database upon acceptance for publication.

## Use of AI

During preparation of this manuscript, the authors used Claude (Anthropic) and ChatGPT (OpenAI) to critique the manuscript for logical clarity, identify inconsistencies in presentation, and assist with language editing. No figure or data were generated by these tools. The authors reviewed and edited all AI-assisted wording and take full responsibility for the content of the published article.

## Figures

**Fig. S1.**
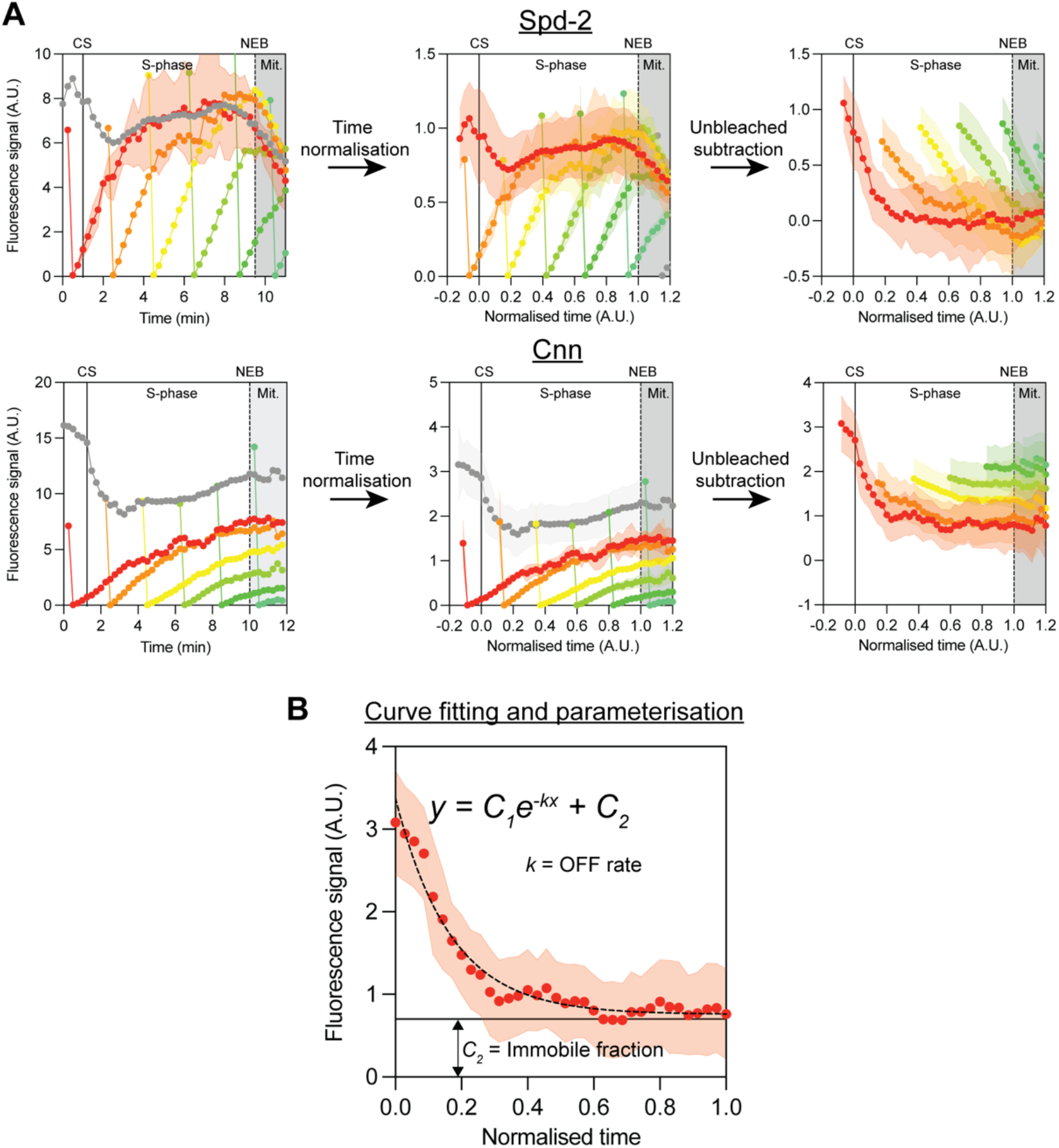
Extracting OFF rates from FRAP data. **(A)** Graphs illustrate the processing steps for Spd-2-NG (top) and NG-Cnn (bottom) FRAP data from a representative embryo. Left graphs show the raw centrosomal fluorescence intensity over real time. Each coloured line represents the mean ± SD fluorescence intensity of 3–4 centrosomes that were bleached at the same time (as in Fig. 2B). The grey line is the mean fluorescence intensity of unbleached centrosomes. Middle graphs show the same data after time normalisation (0 = centrosome separation, solid line; 1 = NEB, dashed line; grey shading, mitosis). This allows the data from embryos with different cycle lengths be more accurately compared. Right graphs show each bleached curve subtracted from the unbleached curve. This removes the change in total centrosomal fluorescence levels generated by the protein binding to or unbinding from the centrosome during the cycle, leaving only the exchange of bleached for unbleached protein to be analysed. **(B)** Schematic of the curve fit. Each difference curve is fitted to an exponential decay to a plateau. The decay constant k gives the OFF rate and the plateau gives the fraction of protein that did not exchange during the time course of fluorescence recovery.

**Fig. S2.**
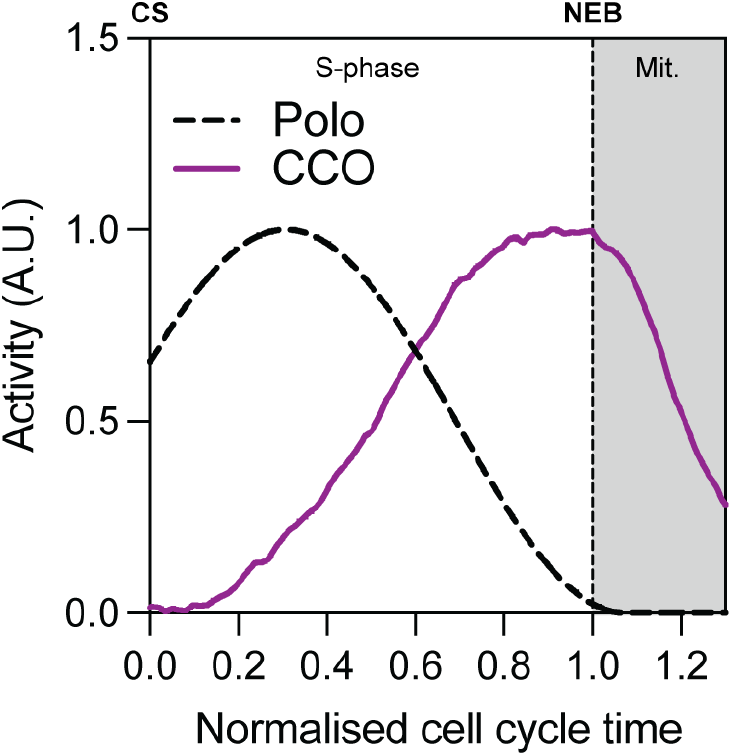
Model inputs: centriolar Polo activity and CCO activity. Graph shows the time course of the two inputs that drive the model (Equations 1 and 2) during NC12. Centriolar Polo activity (P; dashed line) is a model input, not a measurement. It is an offset sinusoidal pulse that approximates the centriolar Polo pulse described previously set to zero (21). CCO activity (F; *purple* line) is the emission ratio of a FRET-based Cdk1 activity sensor (49). It reports the overall balance of cell cycle kinase to phosphatase activity, averaged from N = 6 embryos. Both curves are normalised to a maximum of 1, and F is also rescaled to a minimum of 0. Time is normalised (0 = centrosome separation (CS), 1 = nuclear envelope breakdown (NEB); grey shading, mitosis).

**Fig. S3.**
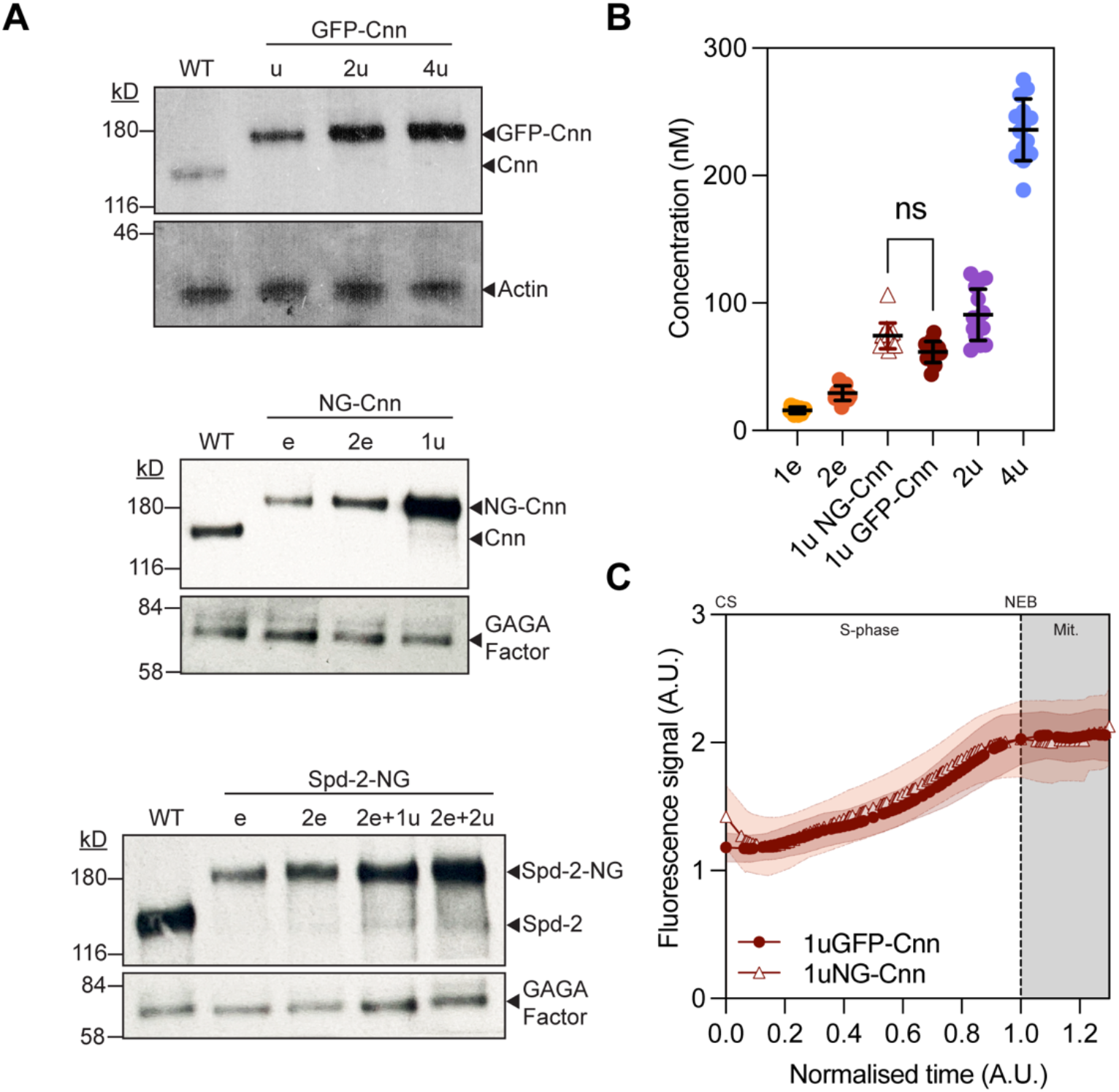
Expression levels of tagged Spd-2 and Cnn lines. **(A)** Western blots of embryo extracts probed with anti-Cnn or anti-Spd-2 antibodies from the GFP-Cnn (top), NG-Cnn (middle) and Spd-2-NG (bottom) lines used for the FCS experiments shown. in Fig. 5. The tagged proteins are expressed in a background that lacks endogenous protein. WT embryos express untagged protein only, to show the endogenous level for comparison. Actin (top) and GAGA factor (middle, bottom) are shown as loading controls. The 2e Cnn line and the 2e+2u Spd-2 line most closely match endogenous levels and so are set to 1 in Fig. 5C. **(B)** Dot plots show the cytoplasmic concentrations of the Cnn lines, measured by FCS at the start of NC12. Each point is one embryo. Bars show mean ± SD. The 1u NG-Cnn and 1u GFP-Cnn lines have similar concentrations (Dunn’s multiple comparisons test after a Kruskal-Wallis test; ns, not significant). **(C)** Mean centrosomal fluorescence of 1u GFP-Cnn and 1u NG-Cnn during NC12, after rescaling. The 1u NG-Cnn trace was scaled to match the starting and plateau intensities of the 1u GFP-Cnn trace. The same factor was applied to the 1e and 2e NG-Cnn data in Fig. 5B,C. Time is normalised (0 = centrosome separation, 1 = NEB; grey shading, mitosis). Shaded bands show SD.

## Supplementary table

**Table S1:** *Drosophila* stocks used in this study.

| Allele | Source |
|---|---|
| *cnn<sup>f04547</sup>* | Exelixis stock no. f04547, Exelixis Stock Centre (Harvard Medical School, Boston, MA). |
| *cnn<sup>HK21</sup>* | (72) |
| *Spd-2<sup>z35711</sup>* | (73) |
| *Spd-2<sup>G20143</sup>* | (59) |
| *polo<sup>10</sup>* | (74) |
| Spd-2-NG | (21) CRISPR Knock-in of Neongreen |
| NG-Cnn | (25) CRISPR Knock-in of Neongreen |
| Ubq-Spd-2-NG | (21) |
| Ubq-NG-Cnn | (21) |
| Ubq-GFP-Cnn | (58) |
| Ubq-Spd-2-GFP | (59) |
| Ubq-Spd-2-CDK20A-NG | (41) |
| Cdk1-FRET | (49) expressed under the maternal α-tubulin promoter; gift from Stefano Di Talia |
| Jupiter-mCherry | Generated by the Daniel St Johnston lab |

**Table S2:** *Drosophila* stocks used in specific experiments.

| Genotype | Purpose | Figure |
|---|---|---|
| Spd-2-NG | FRAP, FCS, Dynamics | 2B, C, D; 3A; 5; 6B; 7; 8 |
| NG-Cnn | FRAP, FCS, Dynamics | 2B, C, D; 3B; 5; 6B |
| Spd-2-NG / *Spd-2<sup>z35711</sup>* | FCS, Dynamics | 5; S3A |
| Ubq-Spd-2-NG / + ; Spd-2-NG | FCS, Dynamics | 5; S3A |
| Ubq-Spd-2-NG; Spd-2-NG | FCS, Dynamics | 5; S3A |
| NG-Cnn / *cnn<sup>f04547</sup>* | FCS, Dynamics | 5; S3 |
| Ubq-GFP-Cnn, *cnn<sup>f04547</sup>*/ *cnn<sup>HK21</sup>* | FCS, Dynamics | 5; S3 |
| Ubq-GFP-Cnn, *cnn<sup>f04547</sup>* | FCS, Dynamics | 5; S3 |
| Ubq-GFP-Cnn, *cnn<sup>f04547</sup>* ; Ubq-GFP-Cnn | FCS, Dynamics | 5; S3 |
| Ubq-NG-Cnn, *cnn<sup>f04547</sup>*/ *cnn<sup>HK21</sup>* | FCS, Dynamics | S3 |
| Ubq-GFP-Cnn, *cnn<sup>f04547</sup>*/ *cnn<sup>HK21</sup>* ; Spd-2-mCherry, *Spd-2<sup>G20143</sup>* / *Spd-2<sup>z35711</sup>* | Dynamics | 6A |
| Ubq-Spd-2-GFP, *Spd-2<sup>G20143</sup>* / *Spd-2<sup>z35711</sup>* | Dynamics (Colchicine or Cyclin RNAi injection) | 7 |
| Ubq-GFP-Cnn, *cnn<sup>f04547</sup>*/ *cnn<sup>HK21</sup>* | Dynamics (Colchicine or Cyclin RNAi injection) | 7 |
| Ubq-Spd-2-CDK20A-NG / + ;; *Spd-2<sup>z35711</sup>* / + | FRAP | 8 |
| *cnn<sup>f04547</sup>* / +; Spd-2-NG | Dynamics, FRAP | 8 |
| w<sup>1118</sup>; Cdk1-FRET ; Jupiter-mCherry | Cdk1 activity (FRET) | S2 |

## Appendices

## Appendix I – FRAP response curve

We estimate loss rates in an evolving system by performing a FRAP analysis. Suppose that the fluorescence response curve of a protein, *y*, satisfies the first-order equation

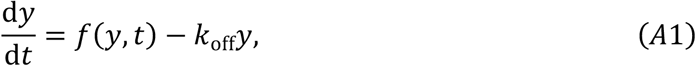

for some unspecified recruitment function *f*. The FRAP response curve of the same protein, photobleached at a time *t* = *t*_0_, is given by

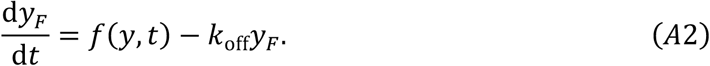

Subtracting equation (A2) from equation (A1) and solving the result gives the exponential decay relationship presented in the main text, subject to the appropriate normalisation. We estimate *k*_off_ by fitting this curve to a FRAP experiment. It is also possible to estimate a varying loss rate provided that *k*_off_ evolves sufficiently slowly (i.e. quasi-statically).

## Appendix II – Derivation of PCM assembly model

We model PCM assembly as a coupled system between Spd2 and Cnn that is driven by two functional inputs; the centriolar Polo activity, *P*(*t*), and the kinase/phosphatase ratio (i.e. the cell cycle oscillator, CCO), *F*(*t*).

### Spd2

We assume that the Spd2 scaffold is generated at the centriole surface, with Polo activity at the centriole acting enzymatically to facilitate the phosphorylation and recruitment of Spd2. We also assume that this phosphorylated Spd2 scaffold can disassemble back into its original cytoplasmic form, and that the rate of this disassembly increases as the CCO-driven kinase/phosphatase ratio increases. We model this by supposing that the disassembly rate of Spd2 is given by *k*_*S*_^−^(1 + *λF*), where *k*_*S*_^−^ is the base disassembly rate, and *λ* is a constant that measures how responsive the disassembly rate is to the CCO.

We denote the cytoplasmic concentration of Spd2 by *S*_Cyt_ and the phosphorylated Spd2 scaffold by *S*_1_. We also distinguish the centriolar Polo that is bound to cytoplasmic Spd2 in the enzyme complex, *S*_Cyt_: *P*^∗^, from the centriolar Polo that is not bound to Spd2, *P*^∗^, and assume that the total centriolar Polo activity is given by *P* = *P*^∗^ + *S*_Cyt_: *P*^∗^. This is our known functional input. With these definitions, the system of reactions reads

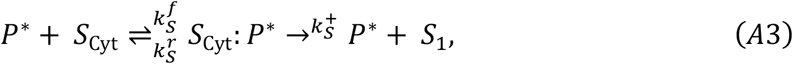

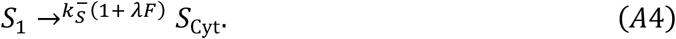

There are several possibilities for how the catalytic conversion reaction shown in equation (A3) may take place. For example, Spd2 may bind to the Polo receptors at the centriole surface—likely Ana1/CEP295 —or it may involve an independent Spd2-receptor, or it may occur away from the centriole surface (i.e. without any Spd2-specific receptors) in a nearby region of high kinase activity. However, these scenarios may be modelled using equivalent mathematics and therefore we do not attempt to distinguish them here.

Next, we suppose that the Cnn scaffold, *C*, stabilises the *S*_1_ scaffold at a rate *k*^*f*^_stab_. We label this stabilised scaffold *S*_2_. We suppose that the rate of disassembly of this scaffold is given by *δk*_*S*_^−^, where *δ* < 1 is the stabilisation factor (i.e., the extent to which the S2 scaffold is more stable than the S1 scaffold). We further suppose that *S*_2_ can disconnect from the Cnn scaffold and destabilise spontaneously back into its *S*_1_ form at a rate *k*^*r*^_stab_. These reactions read

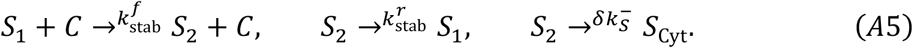

By the law of mass action, we write these reactions as a system of odes, which reads

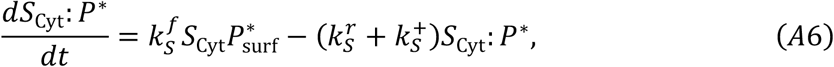

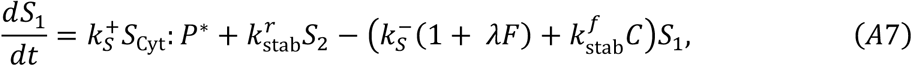

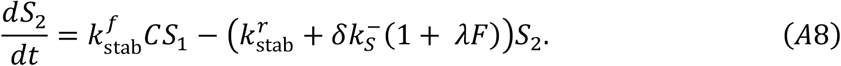

Under the assumption that the production rate is slower than the Polo binding/unbinding rates, 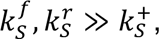 we find that the enzyme reaction may be described by Michaelis-Menten kinetics;

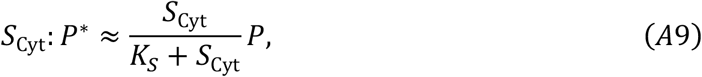

where *K_S_* = *k^r^_S_*/*k^f^_S_*. Similarly, if we assume that the equilibrium between the unstable *S*_1_ scaffold and the stable *S*_2_ scaffold is reached rapidly, 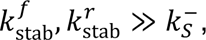 we find that

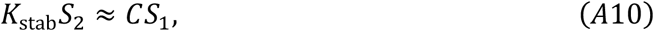

where *K_stab_* = *k^r^_stab_*/*k^f^_stab_*. Summing equations (A7) and (A8), and substituting expressions (A9) and (A10) where applicable, yields the governing equation for the total Spd2 scaffold, *S* = *S*_1_ + *S*_2_;

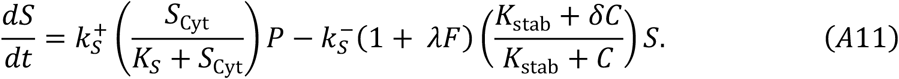

If the rate of turnover is sufficiently rapid relative to the rate of change of the Polo activity, we can approximate equation (A11) using the quasi-static limit which reads

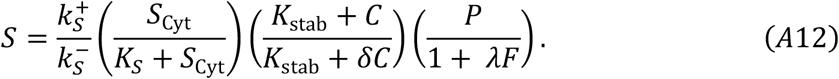

### Cnn

We suppose that Cnn has *N* potential phosphorylation states, and we let *C*_*n*_ denote Cnn with *n* sites that are phosphorylated. We suppose that Spd2 scaffold enzymatically catalyses the production of the Cnn scaffold into its fully-phosphorylated state,

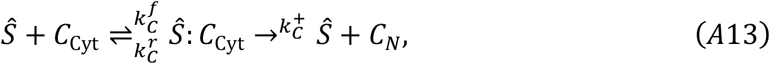

where *S* = *Ŝ* + *Ŝ*: *C*_Cyt_ is the total amount of Spd2 scaffold. Note that we have not distinguished between scaffolds *S*_1_ and *S*_2_ with respect to Cnn production in order to simplify the mathematics.

For simplicity, we suppose that Cnn must be dephosphorylated *N* times, reaching the state *C*_0_, before it can be released from the scaffold at a rate *k*_*S*_^−^. In reality, Cnn may gradually become more prone to be released from the scaffold as it becomes more dephosphorylated, but this simplification captures the relevant biology while considerably simplifying the analysis. We model this by assuming that cytoplasmic phosphatase facilitates dephosphorylation, and that active Polo bound to Spd2 is able to re-phosphorylate sites on the Cnn molecule and strengthen its position. For simplicity, we assume that the amount of active Phosphatase in the cytoplasm is inversely proportional to the time-dependent CCO kinase activity, i.e. Phosphatase ∝ 1/*F*(*t*) and we suppose that phosphorylation by Spd2-bound Polo occurs at a constant rate. With this setup, disassembly of the Cnn scaffold can be modelled with the reaction system

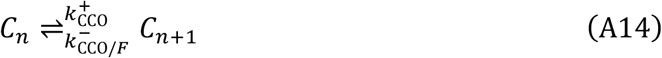

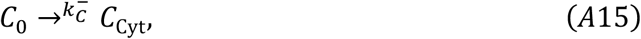

for *n* = 0 ⋯ *N* − 1, with the additional constraint that *C* = ∑ *C*_*n*_. By the law of mass action, reactions (A13) – (A15) yield the system of ode’s,

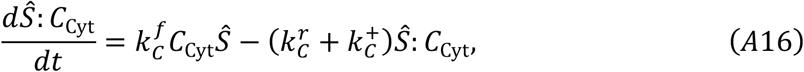

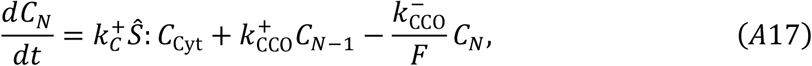

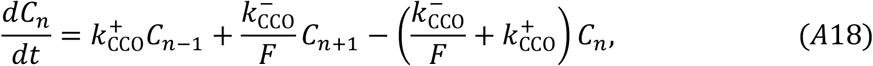

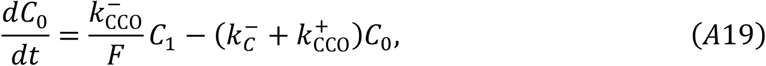

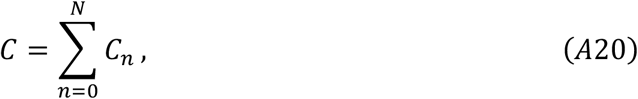

where equation (A18) applies to *n* = 1 ⋯ *N* − 1. As with the Spd2 production model, we assume that the Cnn production rate is slower than the binding/unbinding rates, 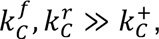 so that the enzyme reaction may be described by Michaelis-Menten kinetics,

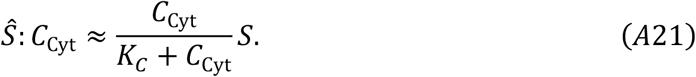

Summing equations (A17) – (A19) and substituting in equation (A21) gives

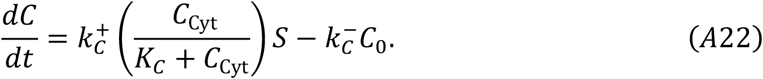

To close the system, we need to write *C*_0_ in terms of *C*. Under the assumption that the rates of phosphorylation and dephosphorylation are sufficiently fast, 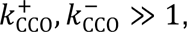 we find from equation that *C*_*n*_ evolves quasi-statically according to the recurrence relation

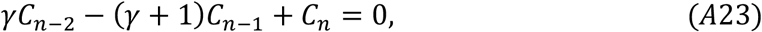

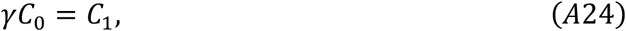

where *γ* = *K*_CCO_*F*, and *K*_CCO_ = *k*^+^_CCO_/*k*^-^_CCO_. Solving this recurrence relation gives

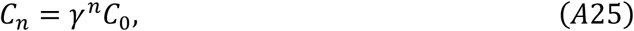

and hence we find that

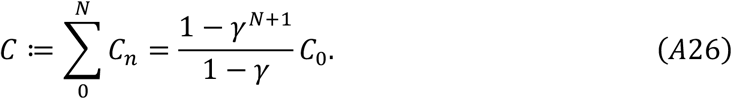

Substituting this result into (A22) gives the governing equation for *C*,

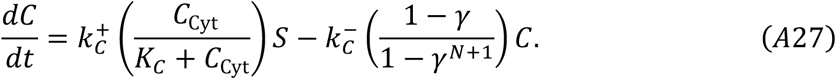

To simplify this expression, we consider the limit *N* → ∞ since there is experimental evidence that the number of phosphorylation sites that bind Cnn to the scaffold is large (∼10). We must consider two cases. First, if *γ* > 1, we observe that

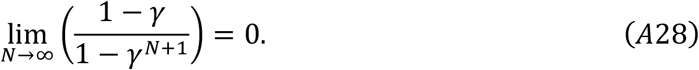

On the other hand, if *γ* < 1, we find that

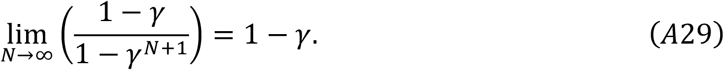

Hence, the evolution of *C* in the large-*N* limit is given by

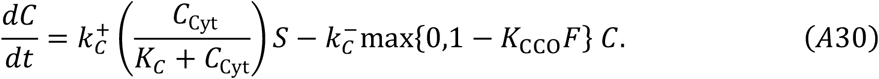

Equations (A11) (or its quasi-static equivalent (A12)) and (A30) together describe the PCM assembly model given an input for the centriolar Polo activity, *P* and CCO, *F*

We assume that the parameters in these equations are non-dimensionalised such that the peak values of *S* and *C* are equal to unity in cycle 12 in the wild type regime. We also suppose that time is non-dimensionalised so that the length of cycle 12 between centrosome separation and nuclear envelope breakdown is equal to unity.

## Appendix III – Estimating model parameters

There are 10 unknown parameters in this model, two recruitment rates from the cytoplasm to the centrosome, *k*^+^_*S,C*_, two loss rates at the centrosome, *k*^-^_*S,C*_, four dissociation constants, *K*_*S*,*C*,stab,CCO_, the stability factor *δ*, and the CCO sensitivity factor, *λ*. In the following, all estimates are given to two significant figures.

### Estimating K_S_ and K_C_

To estimate *K*_*S*_ from experimental data, we assume that, when the Spd2 scaffold is at its largest, the Cnn scaffold is sufficiently large that the Spd2 scaffold is nearly fully saturated with Cnn. Under this assumption, the size of the Spd2 scaffold at its peak is given by considering the quasi-static expression for Spd2, equation (A12), in the limit *K*_stab_ ≪ *δC*. This reads

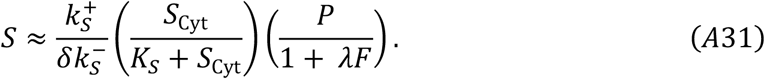

Given that *P* and *F* are functional inputs that do not depend on the amount of Spd2 in the cytoplasm, it follows that the peak size of the Spd2 scaffold scales with the cytoplasmic concentration of Spd2 as *S*_Cyt_/(*K*_*S*_ + *S*_Cyt_). Hence, by fitting *S*_Cyt_/O*K*_*S*_ + *S*_Cyt_P to the peak Spd2 level vs cytoplasmic concentration of Spd2 (Figure 5C), we find that *K*_*S*_ ≈ 1.3.

Similarly, since *S* does not depend strongly on the cytoplasmic concentration of Cnn when the Cnn scaffold is large, it follows from equation (A30) that the peak size of the Cnn scaffold scales with the cytoplasmic concentration of Cnn as *C*_Cyt_/(*K*_*C*_ + *C*_Cyt_).

Hence, by fitting *C*_Cyt_/O*K*_*C*_ + *C*_Cyt_P to the peak Cnn level vs cytoplasmic concentration of Cnn (Figure 5C), we find that *K*_*C*_ ≈ 2.4.

### Estimating *k*_*S*_^−^, *δ*, *λ*, and *K*_stab_

The off rate of Spd2 may be read from equation (A11) as

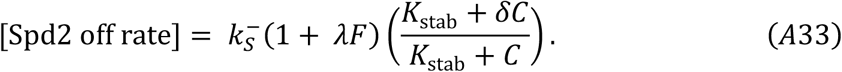

We performed FRAP analysis on Spd2 to estimate the off rate given by equation (A33), which varies over the course of the cycle (Figure 2C). By analysing different parts of the cycle, we are able to estimate the different unknown parameters in this expression.

- *Early cycle.* At the start of the cycle, the Cnn scaffold is small (so we set *C* = 0), and the kinase/phosphatase ratio is low (so we set *F* = 0) and therefore [Spd2 off rate] = *k*_*S*_^−^ ≈ 7.2.
- *Mid cycle.* We suppose the Spd2 scaffold is saturated with Cnn by the mid cycle, and therefore the variation between the off rate at *t* = 0.5 ([Spd2 off rate] ≈ 2.9) and *t* = 1 ([Spd2 off rate] ≈ 4.4) is given by (1 + *λF*(*t* = 0.5))/(1 + *λF*(*t* = 1)) = 2.9/4.4. Evaluation the CCO at these timepoints (*F*(0.5) = 0.47 and *F*(1) = 1) and solving this equation gives *λ* ≈ 1.8.
- *Late cycle.* When the CCO reaches its peak, *F* = 1, the Spd2 scaffold is fully saturated by Cnn and therefore the off rate is given by ([Spd2 off rate] ≈ *k*_*S*_^−^(1 + *λ*)*δ* ≈ 4.4. Substituting in the previously found values gives *δ* ≈ 0.22.
- *Transition.* Finally, we choose *K*_stab_ so that the rate at which the off-rate transitions between the high, unsaturated regime in the early cycle to the low, Cnn-saturated regime in the mid to late cycle is in agreement with the data. We find *K*_stab_ ≈ 0.05 to be an appropriate value.

### Estimating *k*_*C*_^−^ and *K*_CCO_

Performing FRAP analysis on Cnn at multiple times in the cycle demonstrates significant loss at the beginning of the cycle that rapidly reduces to near-zero as the cycle progresses (Figure 2C). Since CCO activity is lowest at the start of the cycle and rapidly increases to a peak as the cycle progresses, our model for the off rate presented in equation (A30), [Cnn off rate] = *k*_*C*_^−^max{0,1 − *K*_CCO_*F*}, is consistent with this data. At the start of the cycle, *F* ≈ 0, and therefore we can directly read that *k*_*C*_^−^ ≈ 5.7 from the initial off rate in the high phosphatase, low kinase regime. Since the off rate of Cnn decreases to zero at approximately *t* = 0.5, we find that *K*_CCO_ ≈ 1/*F*(0.5) ≈ 3.5.

### Estimating *k*_*C*_^+^, *k*_*C*_^+^

We find *k*_*S*_^+^ and *k*_*C*_^+^ by fitting the model with the constraints that Spd2 and Cnn reach a peak value of 1. We find that *k*_*S*_^+^ ≈ 10.8, and *k*_*C*_^+^ ≈ 5.4. For simplicity, we suppose that the PCM starts with zero initial material *S*(*t* = 0) = *C*(*t* = 0) = 0 in these simulations unless otherwise stated.

In Figure 4C, we present the parameters and their dimensionless values as calculated above. It is possible to determine the dimensional values of some of these parameters by expressing them relative to the scalings we used to non-dimensionalise the system. These scalings are the cytoplasmic concentration of Spd2 and Cnn in cycle 12, [*S*_Cyt_] ≈ 209nM and [*C*_Cyt_] ≈ 29nM, and the time between CS and NEB (approximately 8 minutes), [*t*] ≈ 480s.

Unfortunately, the peak concentration of Spd2, Cnn, and Polo molecules at the centrosome in cycle 12, [*S*_WT_], [*C*_WT_], and [*P*_WT_] are not so easy to obtain, since they require knowledge of the absolute number of molecules at the centrosome. For this reason, it is not possible to compute dimensional on rates of scaffold material.

